# Genetic determinants of chromatin reveal prostate cancer risk mediated by context-dependent gene regulation

**DOI:** 10.1101/2021.05.10.443466

**Authors:** Sylvan C. Baca, Cassandra Singler, Soumya Zacharia, Ji-Heui Seo, Tunc Morova, Faraz Hach, Yi Ding, Tommer Schwarz, Chia-Chi Flora Huang, Cynthia Kalita, Stefan Groha, Mark M. Pomerantz, Victoria Wang, Simon Linder, Christopher J. Sweeney, Wilbert Zwart, Nathan A. Lack, Bogdan Pasaniuc, David Y. Takeda, Alexander Gusev, Matthew L. Freedman

**Author notes:** These authors jointly supervised this work (Alexander Gusev, Matthew L. Freedman).

## Abstract

Methods that link genetic variation to steady-state gene expression levels, such as expression quantitative trait loci (eQTLs), are widely used to functionally annotate trait-associated variants, but they are limited in identifying context-dependent effects on transcription. To address this challenge, we developed the cistrome-wide association study (CWAS), a framework for nominating variants that impact traits through their effects on chromatin state. CWAS associates the genetic determinants of cistromes (*e.g.*, the genome-wide profiles of transcription factor binding sites or histone modifications) with traits using summary statistics from genome-wide association studies (GWAS). We performed CWASs of prostate cancer and androgen-related traits, using a reference panel of 307 prostate cistromes from 165 individuals. CWAS nominated susceptibility regulatory elements or androgen receptor (AR) binding sites at 52 out of 98 known prostate cancer GWAS loci and implicated an additional 17 novel loci. We functionally validated a subset of our results using CRISPRi and *in vitro* reporter assays. At 28 of the 52 risk loci, CWAS identified regulatory mechanisms that are not observable via eQTLs, implicating genes with complex or context-specific regulation that are overlooked by current approaches that relying on steady-state transcript measurements. CWAS genes include transcription factors that govern prostate development such as *NKX3-1*, *HOXB13*, *GATA2*, and *KLF5*. Moreover, CWAS boosts discovery power in modestly sized GWAS, identifying novel genetic associations mediated through AR binding for androgen-related phenotypes, including resistance to prostate cancer therapy. CWAS is a powerful and biologically interpretable paradigm for studying variants that influence traits by affecting context-dependent transcriptional regulation.

## Introduction

Genome-wide association studies (GWAS) have identified hundreds of thousands of genetic variants associated with human traits and diseases^1^. The majority of these variants map to regulatory elements and confer risk by affecting transcription of nearby genes^2–8^. Determining how non-coding genetic variants contribute to diseases and complex phenotypes has proven an enormous challenge^9–12^. To address this challenge, large-scale efforts have catalogued many thousands of *cis*-acting expression quantitative trait loci (eQTLs)^13–15^. At these loci, the genotype of a single nucleotide polymorphism (SNP) correlates with steady-state expression of a nearby gene (eGene). eQTLs can identify genes that mediate risk^16–19^ and are present at 40-50% of disease-associated genomic loci by some estimates^15, 20^.

It has become increasingly apparent that the utility of eQTLs for mechanistically characterizing genetic risk variants is limited by several factors. eQTLs that are relevant for complex phenotypes are often context-dependent^13–15^. Such eQTLs are not observable at steady-state in bulk, differentiated tissues, but only in certain cell types, at specific developmental stages, or in response to stimuli^21–26^. Steady-state eQTLs are depleted near genes that are likely to contribute to complex phenotypes, including transcription factors, developmental genes, and highly conserved or essential genes^27^. This finding may reflect purifying selection and regulation by redundant or context-specific enhancers^28^. Consequently, steady-state *cis*-eQTLs explain only 11% of the heritability for an average trait by a recent estimate^27, 29^, or up to 25% when transcription is profiled in disease-relevant tissues^30^.

Many eQTLs influence gene expression through effects on chromatin – for instance, by altering regulatory element activity^31–34^. Increasingly, studies have analyzed the effect of risk-associated genetic variants on chromatin itself, rather than the more distal readout of gene expression^32, 35–37^. Analogous to eQTLs, chromatin QTLs (cQTLs) are SNPs whose genotype correlates with chromatin state, characterized by histone modifications, TF binding, or chromatin accessibility^38–41^. In a complementary manner to cQTLs, allelic imbalance (AI) in epigenomic data – differential representation of heterozygous SNP alleles in sequencing reads – can also identify variants that affect chromatin state^24, 25, 37, 42, 43^. Studying mechanisms of GWAS risk variants through their effects on histone modifications or TF binding (the “cistrome”) is conceptually appealing. This approach captures an immediate downstream consequence of genetic variation on chromatin and could be powerful in cases where complex gene regulation obscures the effects of eQTLs. This approach can provide mechanistic insight into how eQTLs affect gene expression and ultimately phenotypes^32, 35, 36, 38, 44^. cQTLs and AI can implicate specific transcription factors whose cognate DNA binding motif is subject to genetic variation^43^. The use of cQTLs and AI for understanding trait heritability is limited, however, by the lack of (1) large panels of reference epigenomes from relevant tissues and (2) a unified framework for integrating these data into GWAS.

Here, we describe a biologically and statistically principled approach for identifying variants that contribute to phenotypes through effects on the cistrome. We introduce a cistrome-wide association study (CWAS), which identifies the genetic determinants of TF binding and chromatin activity and associates genetically predicted chromatin signal with the trait using GWAS summary statistics. Because the biological unit of association to the trait is the regulatory element, CWAS provides direct mechanistic insights. Our approach conceptually parallels transcription-wide association studies (TWAS), which estimate the *cis*-genetic component of expression for each gene and associate this quantity with a trait.^45^

We performed a CWAS of prostate cancer, one of the most heritable and common cancers^46^. We find that heritable variation in the cistrome of the androgen receptor (AR) – a critical TF in prostate cancer pathogenesis, treatment, and progression – mediates risk at 21% of prostate cancer risk loci. In addition, 45% of prostate cancer risk loci can be explained in part by genetic variation in activity of promoters and enhancers, as measured by H3K27 acetylation (H3K27ac). CWAS excels at annotating disease mechanisms specifically at GWAS risk loci that are difficult to discover through eQTL-based analyses. CWAS implicated prostate developmental genes in prostate cancer risk that lack robust eQTLs likely due to complex regulation and/or context-dependent expression.

### Overview of the methods

We developed a systematic approach that links genetic variation in TF binding or chromatin state to trait heritability (**Fig. 1A**). We leverage the growing number of chromatin immunoprecipitation and DNA sequencing (ChIP-seq) datasets from genetically distinct individuals to create epigenomic reference panels (**Fig. 1B**). A limitation of existing ChIP-seq datasets is that most lack SNP genotypes necessary for studying genetic-epigenetic interactions. We therefore created an approach to impute genotypes from ChIP-seq data with high accuracy^47^ (**Fig. S1** and **Supplementary Note**), enabling us to conduct the largest study to date of cistromes across a non-hematologic tissue. We identify genetic determinants of epigenomic features (e.g., AR binding or H3K27 acetylation) by jointly relating allelic imbalance and peak intensity to the genotypes of nearby SNPs. These models identify SNPs that, alone or in linear combinations, correlate with epigenomic peak intensity. Integrating this information with summary statistics from GWAS, we identify peaks whose genetic determinants are associated with the trait of interest. The result is a cistrome-wide association study (CWAS) that identifies peaks whose genetically predicted activity is associated with risk of a trait or disease (**Fig. 1B**).

**Figure 1.**
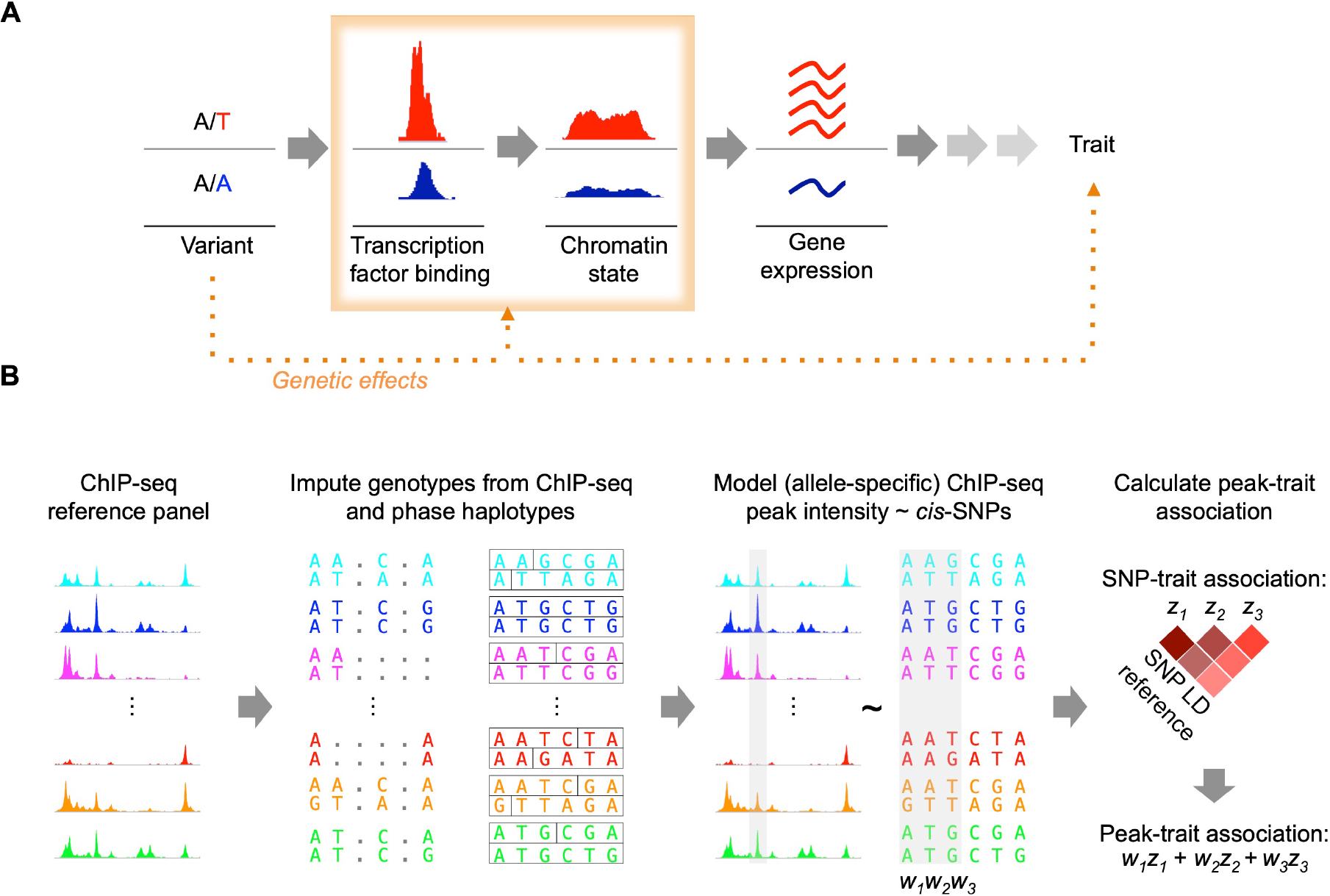
Overview of the method. **(A)** Cistrome-wide association studies identify epigenomic features that are genetically associated with a trait. **(B)** Epigenomic sequencing reads (ChIP-seq and ATAC-seq) are merged on a per-individual basis and used to impute SNP genotypes. Haplotypes are then phased based on reference panels. Normalized read abundance and allele-specific reads at heterozygous SNPs are modeled as a function of *cis*-SNP genotypes. The resulting models capture the genetic determinants of peak intensity.

### cQTLs and allelic imbalance identify tens of thousands of regulatory elements impacted by *cis*-SNPs

We utilized data from two recent studies of prostate cancer epigenomes, which performed ChIP-seq for TFs and histone modifications across a combined 163 individuals of predominantly European ancestry^48, 49^ (**Tables S1-2**, **Fig. S2**). The dataset comprises 131 ChIP-seq experiments for AR and 173 for H3K27ac. Because these samples have not been subjected to genotyping, we used ChIP-seq reads to impute high-accuracy germline genotypes at ∼5.5 million SNPs with a minor allele frequency of ≥ 5%^47, 50^ (**Fig. S1** and **Supplemental Note**).

By analyzing both allelic imbalance (AI) and cQTLs in large epigenomic reference panels, we detected widespread *cis*-genetic regulation of chromatin by common SNPs. A combined test for significant cQTL activity or AI^51^ identified 4,243 AR binding sites (ARBS; 9% of total) and 13,569 H3K27ac peaks (17% of total) where the genotype of nearby SNP (cQTL) correlated with the intensity of a peak (“cPeak”) or was significantly imbalanced in ChIP-seq reads (**Fig. 2A**). AR cQTL activity and AI, which are measured independently, correlated in magnitude and direction (*ρ* = 0.80, *p* < 2.2 × 10^-16^), confirming a shared underlying effect in the population (**Fig. 2B**). Measuring both AI and cQTLs increased the number of peaks under detectable genetic control by roughly over 50% compared to either measure alone (**Fig. 2C**). Genetically-determined H3K27ac peaks overlapped with only 41% of AR peaks (**Fig. 2D**), indicating that TF and H3K27ac ChIP-seq data captured overlapping but distinct genetic regulation. cQTLs overlapped significantly with eQTLs from an independent GTEx study and demonstrated correlated effects on chromatin and gene expression (**Fig. S3** and **Supplemental Note**).

**Figure 2.**
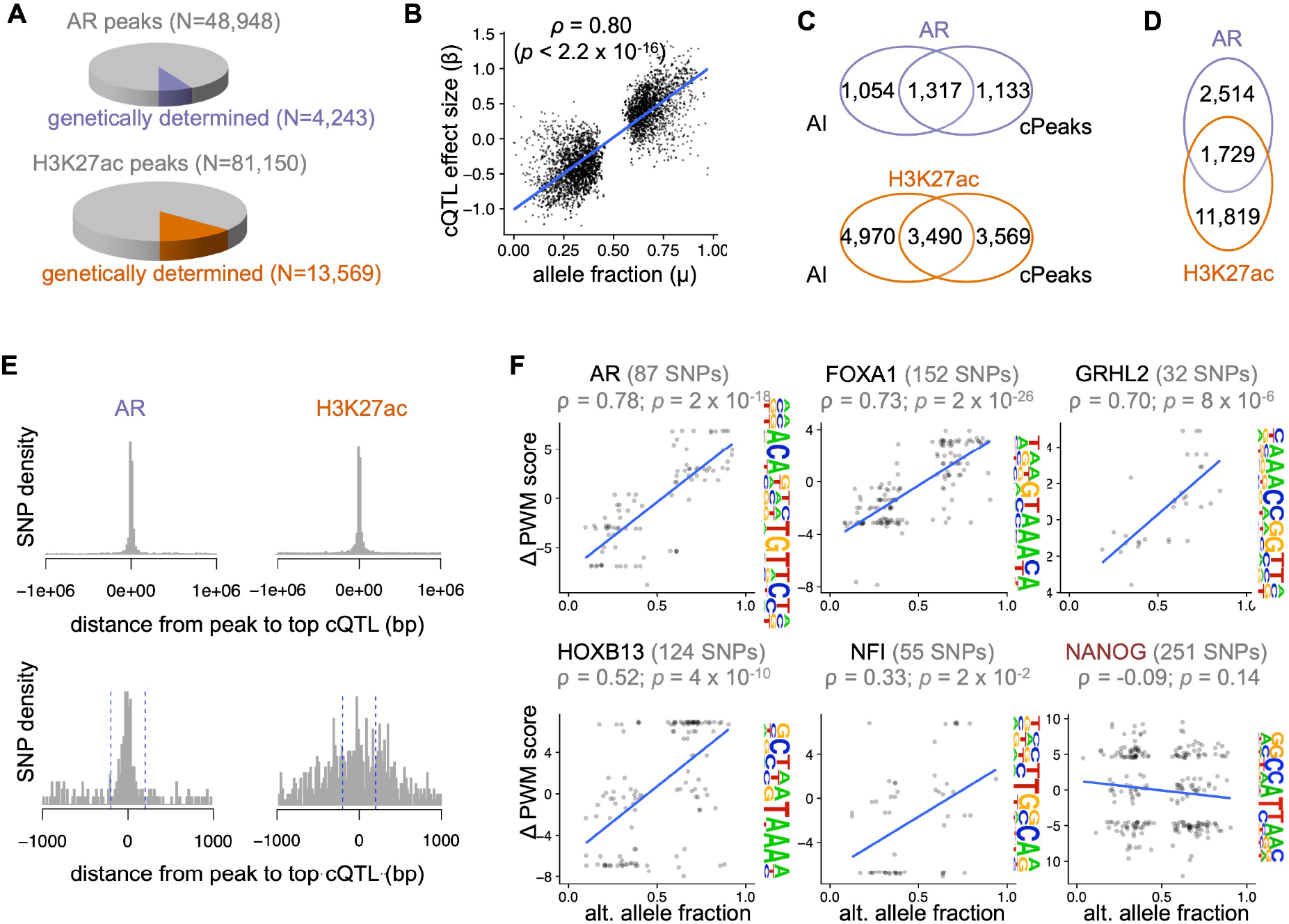
Genetic variation creates abundant chromatin QTLs and allelically imbalanced regulatory elements. (**A**) Portion of all AR and H3K27ac peaks with evidence of genetic determination, defined as a significant combined test for allelic imbalance and cQTL with Q < 0.05 (methods). (**B**) cQTL effect size (*β*) versus allele fraction (*µ*) for peaks with allelic imbalance. *µ* for one SNP per peak is shown. ρ indicates Pearson correlation coefficient. (**C**) Overlap of allelically imbalanced (AI) and chromatin QTL (cQTL) peaks. (**D**) Overlap of genetically determined AR and H3K27ac peaks in (A). (**E**) Distance from the center of significant AR cQTL peaks (permutation-based q value < 0.05) to the corresponding SNP. Blue dashed lines mark ±200bp from the peak center. (**F**) For all heterozygous SNPs overlapping the indicated motif, the difference in the motif position weight matrix (PWM) score for alternate *vs*. reference alleles is plotted against the allele fraction observed in AR ChIP-seq reads. The top five motifs inferred *de novo* from 10,000 randomly selected AR binding peaks are shown. The NANOG motif (red) is included as a negative control.

cQTL SNPs tended to reside in or near peaks: 50% of AR cQTLs and 35% of H3K27ac cQTLs were within 10kb of the corresponding peak center (**Fig. 2E**; **Fig. S4**). Ten percent of AR cQTLs fell within 200 base-pairs of the peak center, suggesting that these SNPs directly affect binding of core TF machinery. Accordingly, 450 heterozygous SNPs within binding motifs of AR and its cofactors demonstrated AI, with AR preferentially binding to the allele that is more similar to the consensus binding motif (**Fig. 2F**), bolstering the functional validity of these QTLs. Nonetheless, 16% of AR cPeaks did not contain a SNP, consistent with distal *cis*-genetic regulation.

### Integrative cistrome models identify genetic determinants of gene regulation

Given the distinct contributions of AI and cQTLs (**Fig. 2C**), we created integrative models combining both features to capture genetic determinants of AR binding and regulatory element activity. We modeled total and allele-specific peak intensity^25, 52^ as a function of all nearby SNP genotypes (**Fig. 3A**). To allow for the possibility that multiple SNPs affect peak intensity, we considered sparse linear models that combine effects from multiple SNPs within 25kb of a peak^45^, an interval that contained 84% of the top 5% of AR cQTLs by significance (**Fig. S4; Supplemental Note**). We refer to these models as “multi-SNP” models. In addition, we included models that incorporate only the SNP most significantly associated with activity (“top SNP” models). Five-fold cross validation demonstrated that 5,580 out of 48,948 AR peaks (11%) and 17,199 out of 81,150 H3K27ac peaks (21%) showed significant correlation between the trained SNP model and peak intensity in held out samples, after correction for multiple hypothesis testing (q < 0.05; **Tables S3-4**).

**Figure 3.**
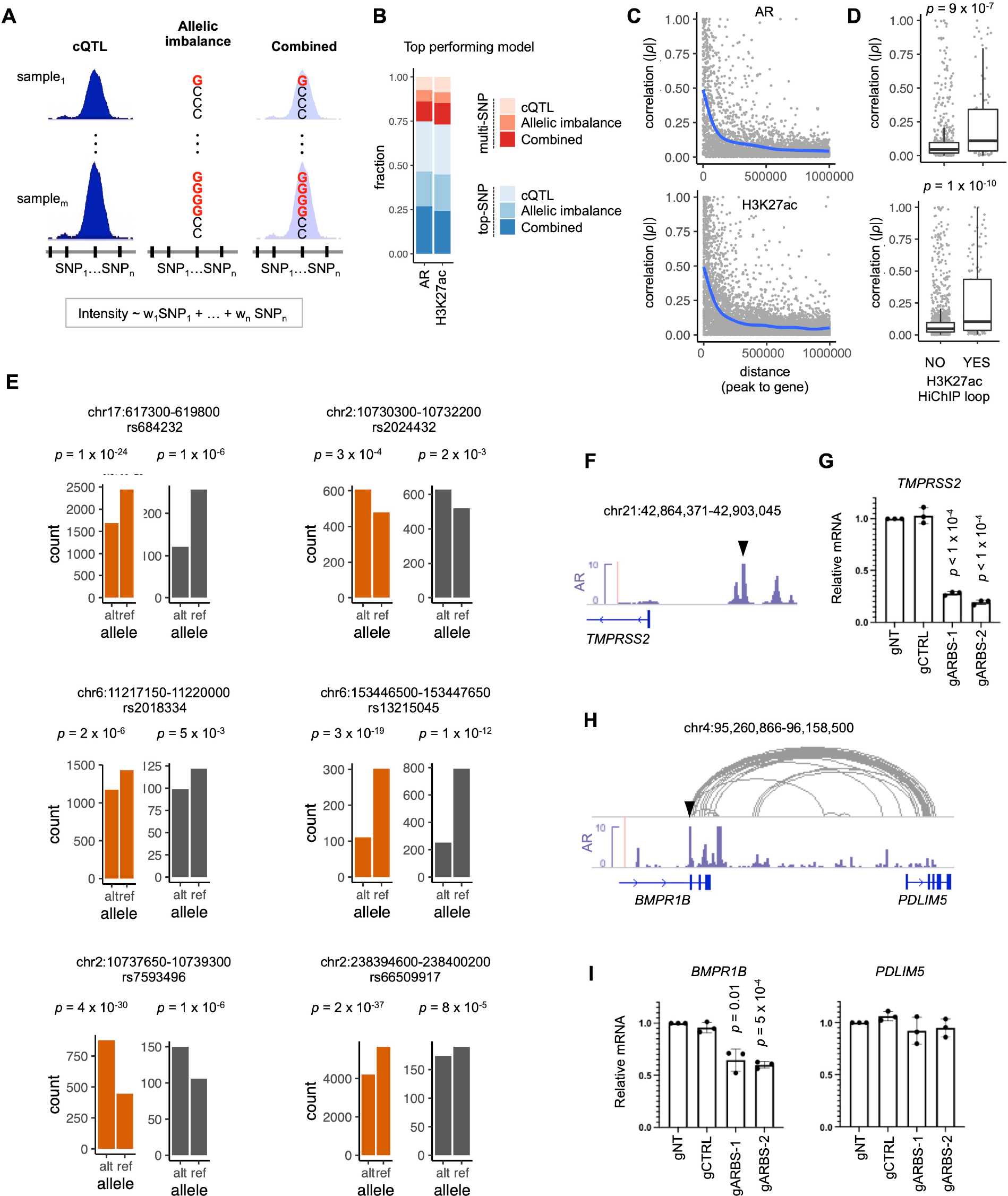
Integrative cistrome models identify genetic determinants of gene regulation. (**A**) Total peak intensity, allele-specific activity, or both are modeled based on *cis*-SNP genotypes. Models include either linear combinations of SNPs (“multi-SNP”), or the single most significantly predictive SNP (“top SNP”; methods). (**B**) Portion of peaks that are best predicted by each model type, based on cross-validation significance. (**C**) Correlation of predicted peak intensity and predicted gene expression. Peak intensity was predicted with total-peak-activity models for individuals in the 1000 Genomes reference panel^85^. Expression of genes within 1Mb was predicted using weights trained on prostate cancer RNAseq data from TCGA^53^ (methods). The absolute value of the Pearson correlation coefficient is indicated as a function of distance between peak and gene. (**D**) AR peak-gene activity correlations shown in (C), stratified by the presence of an H3K27ac HiChIP loop in LNCaP connecting the peak and gene promoter. (**E**) *In vitro* validation of allelically imbalanced regulatory element SNPs. Regulatory elements containing SNPs were assessed for enhancer activity *in vitro* using SNP STARR-seq (Methods). Bar plots indicate reads from reference or alternate haplotypes in H3K27ac ChIP-seq data (orange) and normalized transcript counts for each SNP genotype from SNP STARR-seq (gray). (**F**) Prostate cancer-associated ARBS (black triangle) upstream of *TMPRSS2*. (**G**) Effect on *TMPRSS2* transcript expression with CRISPRi suppression of ARBSs shown in (F). gNT and gCTRL indicate two non-targeting control guide RNAs. (**H**) Prostate cancer-associated ARBS (black triangle) within *BMPR1B*. (**I**) Effect on *BMPR1B* and *PDLIM5* expression with CRISPRi suppression of ARBSs shown in (H).

To assess the genetic architecture of epigenomic features, we compared the performance of “top SNP” and “multi-SNP” models based on cross-validation significance. “Top SNP” models performed the best for the majority of peaks (**Fig. 3B**), indicating the dominant effect of a single SNP. Nonetheless, multi-SNP models outperformed single SNPs for 25% and 27% of AR and H3K27ac peaks, respectively (**Fig. 3B**). This striking finding indicates a simple genetic architecture for most SNP-peak associations: genetic variation in AR binding and regulatory activity often can be ascribed to a single SNP, while multiple SNPs independently affect activity for a minority of peaks.

Our models captured common genetic variation in AR and H3K27ac cistromes that also affect gene expression. We predicted gene expression levels (based on RNA-seq data from prostate cancers profiled in TCGA^53^ and AR and H3K27ac peak intensity using SNPs from 489 genotyped individuals of European ancestry. Across individuals, predicted AR and H3K27ac peak intensity correlated with predicted expression of nearby genes (mean |*ρ*| 0.47 and 0.49 for genes within 10kb for AR and H3K27ac peaks, respectively; **Fig. 3C**). Peak and gene predictions correlated more highly for physically interacting peak-gene pairs than for non-interacting pairs, as assessed by H3K27ac HiChIP in prostate cancer cells^54^ (*p* = 9 × 10 ^-7^ and 1 × 10^-10^ for AR and H3K27ac, respectively; **Fig. 3D**).

We validated allele-specific regulatory activity *in vitro* using an enhancer reporter assay for six H3K27ac peaks (**Fig 3E**; Methods). In addition, suppression of genetically-determined ARBS in LNCaP prostate cancer cells using CRISPRi suppressed the expression of genes linked to these ARBS by H3K27ac HiChIP loops. For instance, suppression of a 14kb-upstream ARBS markedly reduced *TMPRSS2* expression (**Fig 3F-G**; **Table S5**), consistent with a report that this ARBS contains a *TMPRSS2* eQTL^55^. Similarly, suppression of a genetically-determined ARBS decreased expression of its candidate gene based on HiChIP connectivity (BMPR1B; 134KB away) with no effect on the gene containing the ARBS (PDLIM5; **Fig 3H-I**; **Table S5**). These data indicate that our genetic models capture SNPs that influence gene expression through affects on regulatory elements (**Fig. 1A**) and highlight how chromatin conformational data can accurately link cQTL ARBS to the genes they control.

### CWAS identifies prostate cancer risk mediated by genetic variation in AR binding and regulatory element activity

Our genetic models of ARBS and regulatory elements allowed us to identify disease heritability that is likely mediated through effects on these epigenetic features. We performed a cistrome wide association study (CWAS) to associate genetically predicted peak intensity with prostate cancer risk, using summary statistics from a prostate cancer GWAS of 140,306 individuals^56^. Analogous to the framework for a transcriptome wide association study (TWAS)^45^, this approach imputes the genetic component of total and allele-specific peak intensity into populations profiled by GWAS. By utilizing summary statistics from GWAS studies, CWAS takes advantage of the large size of GWAS studies without requiring participant-level information.

CWAS identified 74 ARBs (out of 5,580 ARBS with genetic models) and 199 H3K27ac peaks (out of 17,199) that were significantly associated with prostate cancer risk after Bonferroni correction for multiple hypotheses tested (**Fig. 4A**; **Tables S6-7**). For many CWAS peaks, conditioning the significance of GWAS SNPs on the predicted peak intensity left no significant residual GWAS association, indicating that the genetic contribution of AR binding or regulatory element activity fully accounts for the effect on risk (**Fig. 4B-C** **and Fig. S5A-B**; **Table S6-7**). Conditioning on the CWAS association explained >90% of the GWAS signal for 41% of AR CWAS regions and 52% of H3K27ac CWAS regions (**Table S6-7**). For instance, a single intragenic ARBS within *LMTK2* accounted for the significant GWAS association at this region (**Fig. 4C**). Similarly, H3K27 acetylation at five CWAS peaks near *BIK* and *TTLL12* explained nearby GWAS associations (**Fig. 4C**). In other regions, residual association remained after conditioning on CWAS peaks, suggesting additional mechanisms, more complex regulation, or incomplete tagging^10^ (**Fig. S5C**).

**Figure 4.**
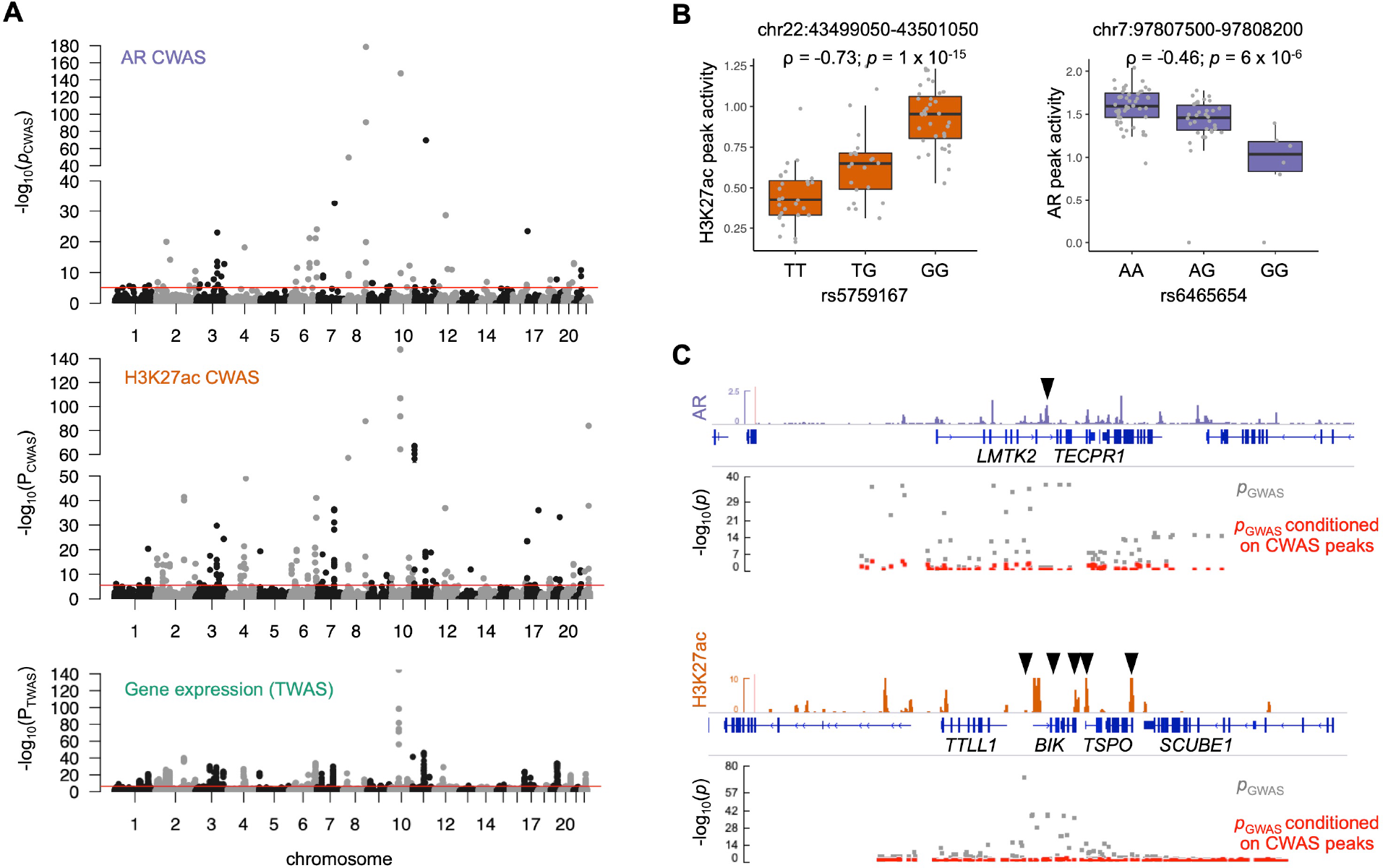
CWAS identifies prostate cancer risk mediated by genetic variation in AR binding and regulatory element activity. (**A**) Manhattan plot showing significant genetic associations with prostate cancer for AR CWAS, H3K27ac CWAS, and TWAS. Red lines indicate genome-wide significance thresholds. (**B**) Normalized read counts at the indicated peaks stratified by genotype of the indicated SNP. (**C**) GWAS SNP significance in the vicinity of the peaks shown in (H), with and without conditioning on genetically predicted activity. The CWAS peaks are marked by a black triangle.

AR and H3K27ac CWAS identified 27 significant “novel” peak-trait associations across 17 regions without a nearby genome-wide significant SNP (**Fig. S6**; **Table S6-7**). CWAS enabled these discoveries mainly by limiting hypothesis testing to SNPs with a high prior likelihood of affecting phenotypes – *i.e.*, testing tens of thousands of genetically determined epigenomic features, as opposed to millions of unselected SNPs. Tested peaks are expected to be enriched for true positive associations, given that prostate cancer risk variants were highly enriched in cQTL ARBS and regulatory elements (**Fig. S7**). In addition, some models achieved higher significance than any single nearby SNP by combining multiple SNPs – for instance, for H3K27ac CWAS peaks near *NEU1* and *BIK* (**Fig. S6B**; **Table S7**). Importantly, novel prostate cancer associations identified by CWAS were subsequently confirmed in a larger GWAS study. After this manuscript was prepared, a prostate cancer GWAS meta-analysis that added ∼94,000 individuals to the GWAS used here^56^ identified novel genome-wide significant associations at 12 of the 17 CWAS-positive GWAS-negative regions^57^.

### CWAS identifies associations not marked by a steady-state eQTL

CWAS uncovered many chromatin-prostate cancer risk associations at eQTL-negative loci, where genetic effects on steady-state gene expression are not observed. We compared CWAS associations to results from TWAS (an integrative analysis of eQTL-trait associations) that used reference gene expression data from 45 tissues (4,448 individuals) including benign prostate tissue and prostate cancer^45, 53^. Some CWAS peaks colocalized with genes identified by TWAS, such as *MLPH* and *MSMB/NCOA4*^53, 55, 58^, but many did not. To compare the relative contributions of TWAS and CWAS in accounting for GWAS risk loci, we defined a set of high-confidence TWAS and CWAS hits where the standardized effect size Z^2^ is greater than 90% of Z^2^ for the top GWAS SNP (i.e. CWAS/TWAS explained >90% of the variance of the top GWAS SNP). Of the high-confidence CWAS associations, 21 ARBS (47%) and 65 H3K27ac peaks (55%) did not map within 1Mb of a high-confidence TWAS gene in any tissue (N=844) (**Table S6-7**).

Compared to TWAS, CWAS nearly doubled the number of GWAS risk loci that could be annotated with plausible risk mechanisms. We defined 98 prostate cancer risk regions by merging ±1Mb windows centered on genome-wide significant SNPs. Of these regions, 52 (53%) contained a high confidence AR or H3K27ac CWAS peak (N=21 and N=44, respectively) compared to 34 (35%) that contained a TWAS gene (**Fig. 5A**). Critically, at 28 regions (29%), CWAS detected a high-confidence peak association in the absence of a high-confidence TWAS gene association. Thus, CWAS implicated regulatory elements at the majority of prostate cancer GWAS risk regions, including many regions that lacked a robust association with steady-state gene expression.

**Figure 5.**
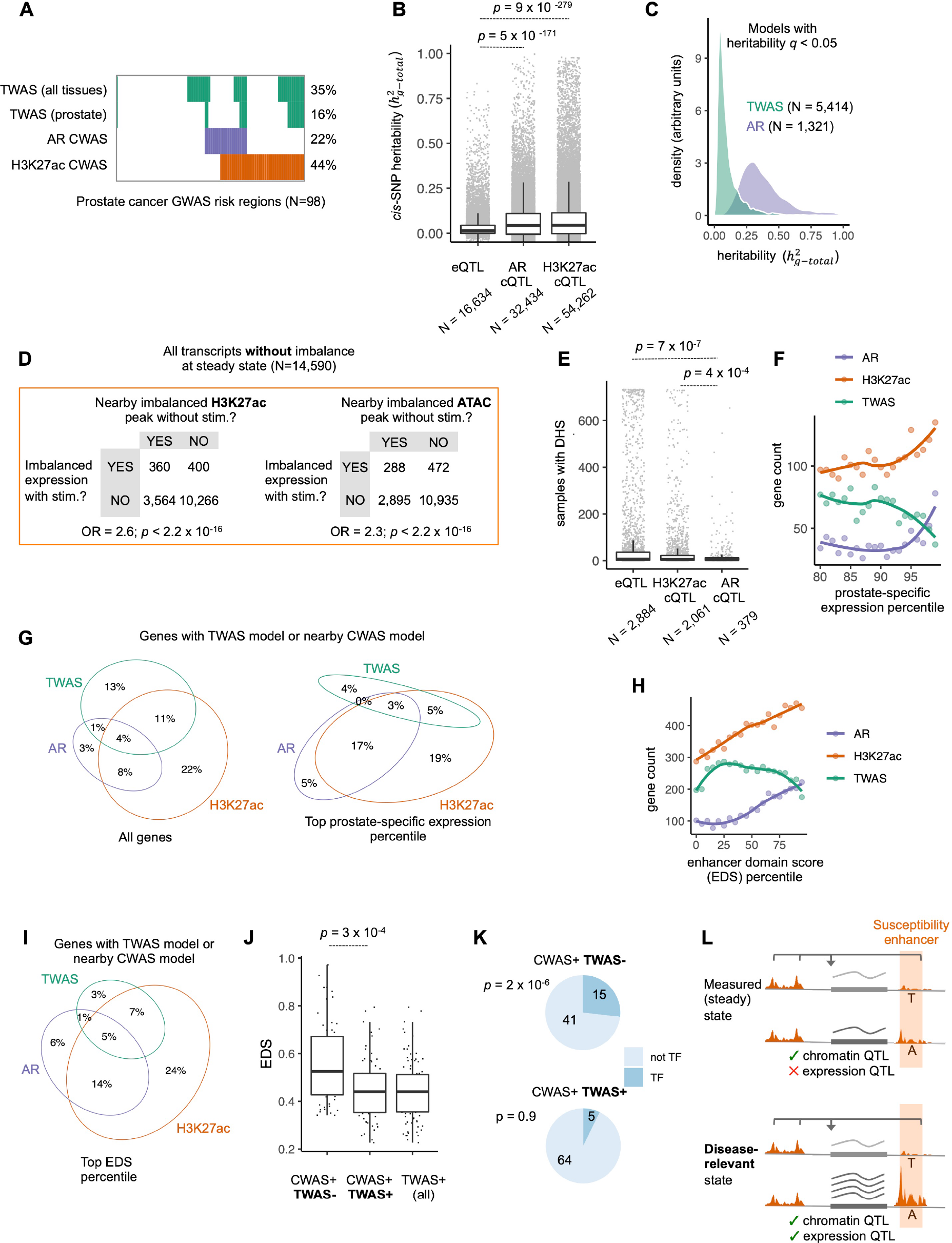
CWAS identifies associations not marked by a steady-state eQTL. (**A**) GWAS risk loci annotated by whether they overlap with a high-confidence CWAS or TWAS peak. TWAS results using reference panels with only prostate tissue or all tissues are shown separately. (**B**) Estimated *cis*-SNP heritability for all assessable genes, AR peaks, or H3K27ac peaks. (**C**) Distribution of heritability estimates for genes or AR peaks with significant heritability (q<0.05). (**D**) Steady-state chromatin measurements revealing context-dependent genetic effects on gene regulation. H3K27ac ChIP-seq, ATAC-seq, and RNA-seq data from LNCaP were generated at baseline and after 16 hours of stimulation with dihydrotestosterone (DHT) and assessed for allelic imbalance^42^. Contingency tables show all transcripts that do not exhibit allelically imbalanced expression at baseline, stratified by (1) whether they demonstrate imbalanced expression with DHT treatment and (2) whether they are within 100kb of an ATAC-seq or H3K27ac peak with allelic imbalance at baseline. Odds ratio (OR) that a transcript with stimulation-induced imbalance falls within 100kb of a peak that is imbalanced at baseline, compared to transcripts without stimulation-induced imbalance. *p*-values from chi-square tests are indicated. (**E**) Number of ENCODE samples (N=733, representing 438 cell types/states)^62^ with DNAse hypersensitivity at cQTL and eQTL SNPs. p-values for Wilcoxon rank-sum tests are indicated. (**F**) Number of genes with a TWAS model or AR/H3K27ac CWAS model (within 100kb) as a function of prostate-specific expression. Expression in prostate was compared to mean across all GTEx tissues to obtain a z-scores, which were binned by percentiles. (**G**) Percent of genes with TWAS models or CWAS models (within 100kb) for all genes (left) and the top percentile of prostate-specific expression (right). (**H**) Data from (F) grouped by enhancer domain score (EDS) percentile. (**I**) Percent of genes with TWAS models or nearby CWAS models for genes in the top EDS percentile. (**J**) Boxplots of EDS scores for genes within central 100kb of the indicated category of GWAS risk regions. (**K**) Number of genes in indicated category of GWAS risk regions that encode TFs. (**L**) Model demonstrating how latent eQTLs are observable as steady-state cQTLs. All p-values indicate Wilcoxon rank-sum tests.

### Prostate cancer cistromes are more heritable than transcriptomes

We considered why CWAS detected chromatin-prostate cancer associations in TWAS-negative (TWAS-) regions despite using substantially smaller reference panels than TWAS. A potential reason is that genetic variation affects the steady-state cistrome more consistently and predictably than it affects transcription: in many instances, SNP genotypes at risk loci correlated more robustly with regulatory element activity than with gene expression. Two examples are androgen-responsive enhancers that regulate *TMPRSS2* and *NKX3-1*^59, 60^. A variant in the *TMPRSS2* enhancer^59^, rs8134657, correlated strongly with AR peak intensity (ρ = -0.52; *p* = 2.8 × 10^-7^; **Fig. S8A**), but only marginally with steady-state *TMPRSS2* expression in prostate cancer (ρ=-0.24; *p* = 0.03). Similarly, rs1160267, which maps within a 6.7 kb-downstream enhancer of *NKX3-1*^60^ and is the most signficant GWAS SNP in the region, correlated with *NKX3-1* enhancer H3K27ac activity (ρ = 0.35; *p* = 1 × 10^-3^) but not with *NKX3-1* expression (**Fig. S8B**).

In addition, steady-state epigenomic features were substantially more heritable than gene expression levels. *cis*-SNP heritability imposes an upper boundary for predictive accuracy for both TWAS and CWAS models. For instance, the *NKX3-1* and *TMPRSS2* enhancers displayed greater heritability attributable to *cis*-SNPs^45, 61^ than the genes they control (**Fig. S8C**). Globally, *cis*-SNPs explained a significantly greater portion of the heritability of AR and H3K27ac total peak intensity 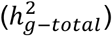 than the heritability of gene expression levels (*p* = 5 × 10^-171^ and *p* = 9 × 10^-279^ for AR and H3K27ac, respectively; **Fig. 5B**). Significantly heritable ARBS demonstrated a median heritability of 0.33 compared to a median heritability of 0.07 for significantly heritable gene expression (**Fig. 5C**). Thus, SNPs explain a greater proportion of variation in regulatory element activity than they explain variation in gene expression levels. This finding supports the hypothesis of lower context-dependent variance for chromatin activity compared to gene expression, leading to more accurate predictive models.

### Steady-state chromatin measurements reveal context-dependent genetic effects on gene regulation

An additional explanation for CWAS associations at TWAS-loci is that steady-state chromatin measurements capture context-dependent genetic determinants of transcription. To test this hypothesis, we measured allelic imbalance in chromatin accessibility (ATAC-seq), H3K27ac, and gene expression data in LNCaP cells at baseline and after 16h of androgen stimulation. We identified 760 transcripts that demonstrated imbalance with stimulation but not at baseline (**Fig. 5D**). These genes were enriched for nearby H3K27ac and ATAC-seq peaks with imbalance in the absence of stimulation (OR 2.3 and 2.6 for ATAC-seq and H3K27ac, respectively, p < 2.2 × 10^-16^; **Fig. 5D**). Thus, effects on expression that are only apparent with stimulation are preceded by genetic effects on nearby regulatory elements at steady-state, as observed previously in immune cells^41^.

Several additional observations support this conclusion. First, tissue- and context-dependent regulatory elements were enriched for steady-state cQTLs compared to eQTLs. We identified eQTLs across tissues profiled in GTEx and cQTLs that overlap with accessible chromatin across 733 tissue samples representing 438 cell types and states^62^. eQTLs tended to localize to chromatin that is accessible in multiple tissues and conditions, while AR and H3K27ac cQTLs overlapped chromatin with more context- or tissue-restricted accessibility (*p* = 7 × 10^-7^ and 4 × 10^-4^, for eQTLs vs. AR and H3K27ac cQTLs, respectively; **Fig. 5E**).

Second, for many genes with prostate-restricted expression (quantified by the z-score for expression in prostate compared to all other tissues), *cis*-SNPs did not correlate with transcript levels but robustly correlated with activity of nearby regulatory elements. We binned genes by quantiles of prostate-specific expression^14^. Then, for each bin we counted genes with a TWAS model (in prostate tissue or prostate cancer) and genes with a nearby CWAS model. Strikingly, genes with increasingly prostate-enriched expression – where power to detect eQTLs should be high due to higher expression levels – were less likely to be modeled by TWAS, but more likely to harbor nearby ARBS or regulatory elements with CWAS models (**Fig. 5F**). While 29% of all genes had a TWAS model in prostate tissues, only 12% of genes in the top percentile of prostate-specific expression had TWAS models. Strikingly, 49% of genes in this top percentile harbored nearby CWAS peaks (**Fig. 5G**), similar to the set of all genes (49%). This top percentile included TWAS-/CWAS+ genes with androgen-responsive expression (e.g., *ACPP* and *SPDEF*) and roles in prostate development (e.g., *FOXA1* and *NKX3-1*).

Third, consistent with prior work^57^, we found that TWAS models were depleted among genes with the highest degree of regulation, as assessed by the enhancer domain score (EDS; **Fig. 5H-I**). In contrast, high-EDS genes were the most likely to have nearby CWAS models (**Fig. 5H-I**). A known limitation of steady-state eQTLs is that they are depleted around highly regulated (high-EDS) genes, which include TFs, developmental genes, and genes involved in disease pathogenesis^27, 28^. This principle may explain the ability of CWAS to annotate prostate cancer risk in TWAS-regions. Prostate cancer risk regions with a CWAS association but no TWAS association (CWAS+/TWAS-) had significantly higher EDS scores than CWAS+/TWAS+ regions (**Fig. 5J**), suggesting that these regions were not captured by TWAS due to more complex regulation and/or correlated factors. CWAS+/TWAS-regions were enriched for TF genes, which are depleted for eQTLs^44^, and contained key prostate developmental genes such as *NKX3-1*, *KLF5*, and *HOXB13* (**Fig. 5K**). Collectively, these results support a model where disease risk is mediated by context-dependent eQTLs that are not observable from steady-state expression, but can be identified in steady-state chromatin (**Fig. 5L**).

### CWAS implicates prostate developmental genes and proto-oncogenes in prostate cancer heritability

The advantages of chromatin models described above allowed CWAS to implicate genes involved in prostate development and oncogenesis that have not been mechanistically tied to prostate cancer GWAS associations. Several such genes, including *MYC*^63^, *KLF5*^64^, *NKX3-1*^65^, *CCND1*^66^, *HOXB13*^67^, and *GATA2*^68^, physically interacted with CWAS ARBS and/or H3K27ac peaks, as assessed by H3K27ac Hi-ChIP (**Fig. 6A-F**). Conditioning GWAS SNP associations upon the genetically predicted peak intensity left little or no residual GWAS significance in these regions, suggesting that regulatory element activity accounts for prostate cancer heritability at these sites.

**Figure 6.**
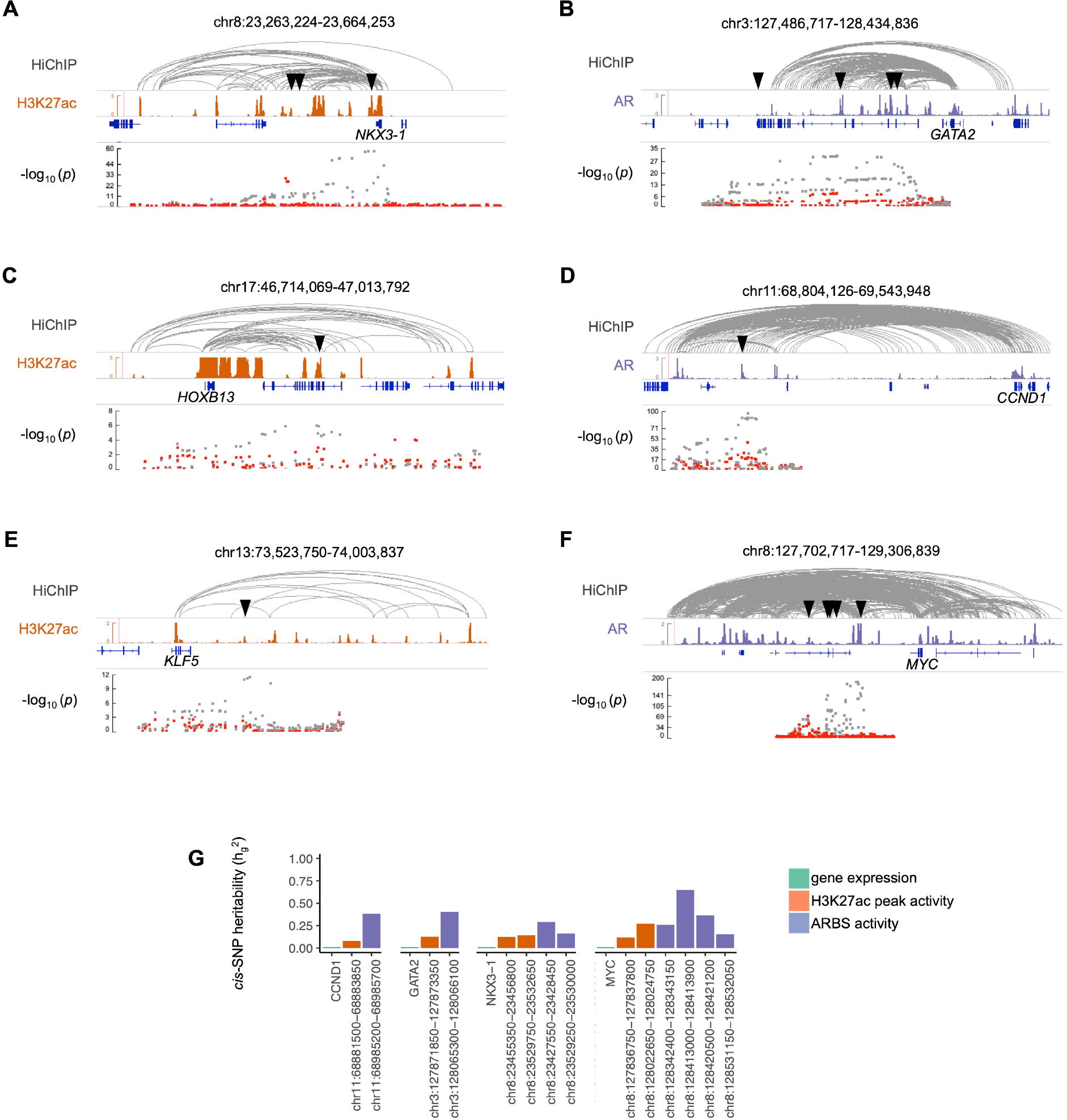
CWAS associations linked to selected prostate developmental genes and proto- oncogenes. **(A-F)** Panels show the (epi)genomic context for CWAS ARBS or H3K27ac for NKX3-1 (**A**), GATA2 (**B**), HOXB13 (**C**), CCND1 (**D**), KLF5 (**E**), and MYC (**F**). For each panel, tracks from top to bottom show H3K27ac HiChIP loops in LNCaP (gray), Normalized read counts for H3K27ac (orange) or AR (purple) ChIP-seq in LNCaP, gene annotations, and significant CWAS H3K27ac peaks or CWAS ARBS (indicated by black triangles). The bottom track shows prostate cancer GWAS SNP significance in the vicinity of the CWAS peaks in gray, and the residual significance after conditioning upon the CWAS H3K27ac peak or ARBS in red. (**G**) *cis*-SNP heritability of indicated genes and CWAS peaks within the regions shown in A-F. Only CWAS peaks with significant *cis*-SNP heritability (*p* < 0.05) are shown.

Importantly, the above genes have not been directly tied to prostate cancer heritability because they lack strong eQTLs and significant TWAS associations. These genes demonstrated low *cis*-SNP heritability of steady-state expression measurements, a likely reason they were not detected by TWAS (**Fig. 6G**). In contrast to gene expression, several peaks associated with the genes above were highly heritable with respect to *cis*-SNPs (**Fig. 6G**). Notably, disruption of the CWAS ARBS ∼220kb centromeric to *MYC* containing the variant rs11986220 was recently shown to impair *MYC* expression, proliferation, and tumorogenesis in a cell-line dependent manner. This finding supports the hypothesis that this ARBS contributes to prostate cancer risk. Thus, by linking epigenomic features to traits, CWAS implicated biologically plausible prostate developmental genes and proto-oncogenes that have been overlooked by analyses based on steady-state expression.

### AR CWAS identifies genetic mediators of diverse androgen-driven phenotypes

We assessed the specificity of AR CWAS by testing the association of AR peak intensity with GWAS risk for additional phenotypes. The median normalized Z^2^ for trait association across AR peaks was greatest for prostate cancer and hypertension, followed by cardiovascular disease and serum testosterone, all of which have known biological links to androgen signaling^69, 70^ (**Fig. 7A**). In contrast, traits without clear links to AR such as asthma or psoriasis showed limited evidence of associations. This finding supports the plausibility that AR CWAS loci act mechanistically through AR or its cofactors.

**Figure 7.**
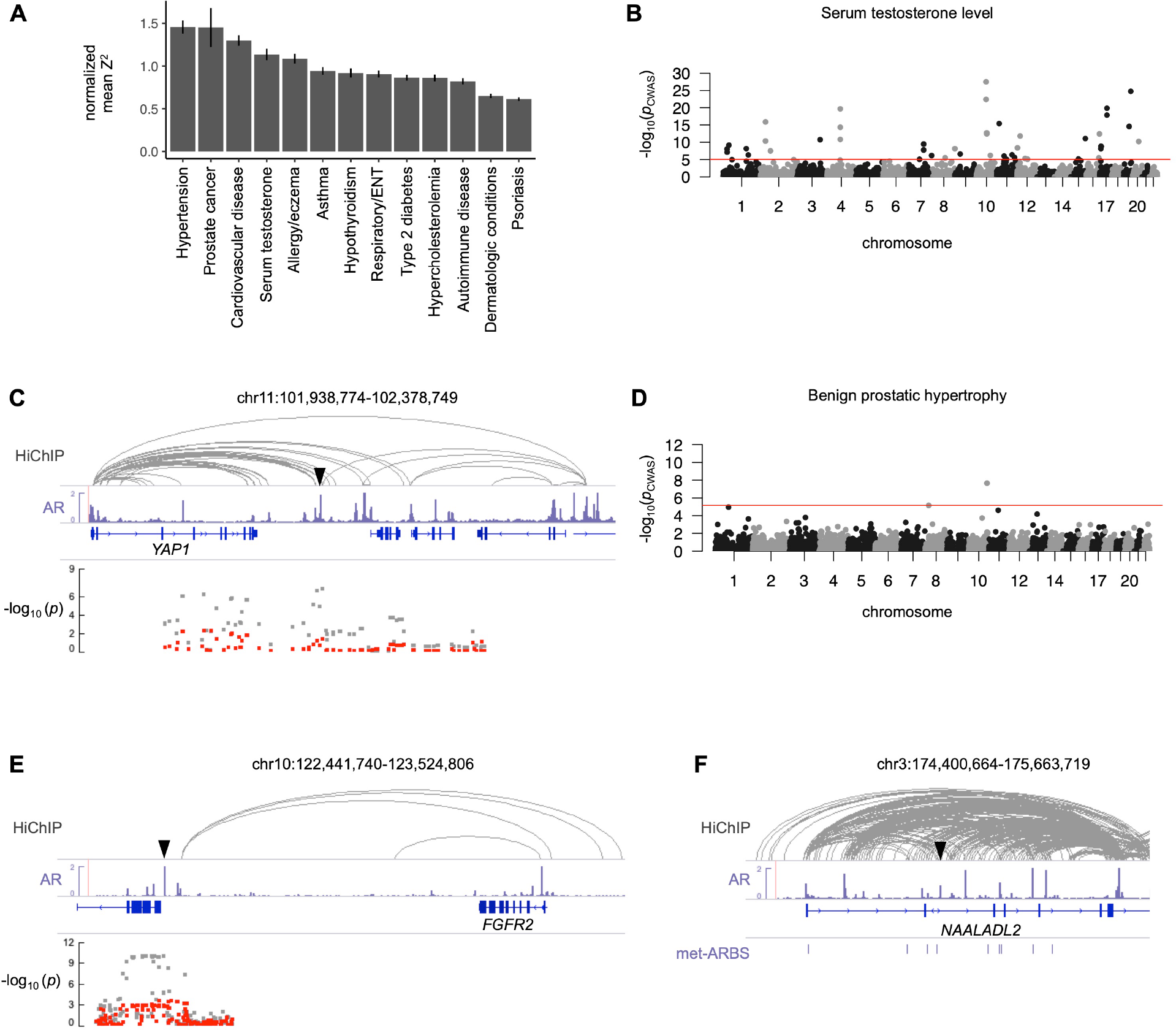
CWAS identifies ARBS underlying heritability of multiple androgen-regulated phenotypes. (**A**) AR CWAS was performed on GWAS for the indicated traits. The mean of the squared effect size (Z^2^) was calculated for each, and normalized to the smallest mean Z^2^ across datasets. Error bars indicate standard error of the mean. (**B**) Manhattan plot showing significance of ARBS associations with testosterone levels among individuals in the UK Biobank^71^. (**C**) Epigenomic context of a significant CWAS ARBS for testosterone near *YAP1.* Tracks from top to bottom show H3K27ac HiChIP loops in LNCaP (gray), normalized AR ChIP-seq read counts in LNCaP (purple), gene annotations, and the location of the significant CWAS ARBS (black triangle). The bottom track shows testosterone GWAS SNP significance in the vicinity of the CWAS peaks in gray and the residual significance after conditioning upon predicted activity of the ARBS in red. (**D**) Manhattan plot showing significance of ARBS associations with BPH among individuals in the UK Biobank. (**E**) Epigenomic context of a significant CWAS ARBS for BPH near *FGFR2*. Tracks are as described for (B). (**F**) Epigenomic context of CWAS ARBS within *NAALADL2* associated with response to androgen deprivation therapy among men with prostate cancer from a clinical trial^77^. Met-ARBs (purple) signify AR binding sites that are enriched in metastatic castration-resistant prostate cancer compared to prostate-localized tumors^49^.

We hypothesized that CWAS could improve discovery power for GWAS of additional AR-mediated phenotypes by focusing on SNPs that genetically influence AR binding. We therefore employed CWAS to identify AR cistrome-mediated heritability of other androgen-associated traits. We identified known and novel regions (N=45) associated with testosterone levels among male UK Biobank participants^71^ (**Fig. 7B**; **Table S8**). The most significant (*p* = 3 × 10^-28^) is an ARBS that contacts *JMJD1C*, a gene with roles in testis development and steroid hormone metabolism^72^ that has been associated with testosterone levels in prior GWAS^73^.

Additional CWAS ARBS interacted with genes implicated by GWAS, including *SHBG*, which encodes sex hormone binding globulin. For 7 of these peaks (16%) a significant GWAS association was not detectable within 1Mb. These novel hits included an intergenic ARBS contacting the *YAP1* promoter (**Fig. 7C**). Knockout of the *YAP1* ortholog in male mice causes degeneration of the adrenal cortex and impairs steroid hormone biosynthesis^74^, supporting the plausibility of this association.

Separately, CWAS of benign prostate hypertrophy (BPH) – another androgen-mediated disease – identified two ARBS associated with this disease (**Fig. 7D**; **Table S9**). The most significantly associated ARBS (*p* = 2 × 10^-8^) was in an intergenic region that physically interacts with the *FGFR2* promoter in LNCaP cells (**Fig. 7E**) and benign prostate tissue^75^. *FGFR2* encodes a receptor highly expressed in prostate stroma that is implicated in the development of BPH^76^. The other BPH-associated ARBS had no genome-wide significant GWAS SNP within 1Mb and localized to an intergenic enhancer of the prostate lineage TF gene *NKX3-1*^49^, which has not been implicated in BPH previously. These results demonstrate that CWAS identifies ARBS accounting for heritability of androgen-related phenotypes at known and novel risk loci.

We reasoned that the enhanced statistical power of CWAS would enable the study of heritability among small populations that are inadequately powered for GWAS. To this end, we applied CWAS to identify genetic determinants of response to androgen deprivation therapy (ADT) among 687 patients with metastatic prostate cancer^77, 78^. No SNPs were associated with ADT response by GWAS at genome-wide significance threshold of p < 5 × 10^-8^. To increase power, we applied CWAS to regions within 1Mb of the 200 most significant SNPs, with Bonferroni correction for 475 tested ARBS. This hypothesis-generating approach nominated an ARBS in intron 2 of *NAALADL2* that was significantly associated with time to progression on ADT (*p* = 7.8 × 10^-5^; hazard ratio 1.29; 95% CI 1.13 – 1.46) **Fig. 7F**; **Table S10)**. *NAALADL2* encodes a glutamate carboxypeptidase that is closely related to PSMA, a therapeutic target in prostate cancer. Expression of *NAALADL2* has been associated with increased grade and stage of prostate cancer, as well as earlier recurrence^79, 80^ and is required for prostate cancer invasion and migration^79^. *NAALADL2* undergoes somatic amplification and rearrangements in 8% of prostate cancers^81^. Additionally, the CWAS ARBS within *NAALADL2* localizes near ARBS that are enriched in metastatic treatment resistant prostate cancer compared to primary prostate cancer^49^ (**Fig. 7F**). Notably, a prior GWAS of prostate cancer aggressiveness required 12,518 prostate cancer cases to identify an association at this gene (*p* = 4.18x10^-8^) meeting genome-wide significance^80^. This finding highlights the power of CWAS for studying therapeutic resistance and other features of interest in small but well-annotated groups such as clinical trial cohorts.

## Discussion

We present the cistrome wide association study (CWAS), a principled and statistically powerful approach for associating genetic variation in regulatory element activity with trait heritability. CWAS provides mechanistic insight into complex phenotypes driven by transcriptional regulation. Applying CWAS to prostate cancer revealed the widespread role of AR in mediating heritability of this disease, which has not been comprehensively characterized until now. CWAS implicated AR binding in 21% of all prostate cancer GWAS risk regions and regulatory element activity in an additional 32%, adding substantially to the number of prostate cancer risk loci that are annotated with plausible mechanisms. Genetic variation in one or a few ARBS accounted for prostate cancer risk at many loci identified by GWAS, such as regulatory elements near *MYC, TMPRSS2*, *GATA2*, and *NKX3-1*. The ability to link these regulatory regions to genes with chromatin conformational data^54, 82^ allows CWAS peaks to be assigned to candidate genes that mediate downstream effects. AR CWAS also detected ARBS at known and novel loci that confer heritability of other AR-mediated traits. We associated AR binding at an enhancer of *NKX3-1* with risk of BPH and a regulatory element near *YAP1* with serum testosterone levels. We experimentally validated the predicted effect of cQTLs on gene expression for six regulatory elements and demonstrated that CWAS ARBS regulate candidate prostate cancer risk genes *TMPRSS2* and *BMPR1B*.

Prior studies have primarily used cQTLs and AI to annotate risk SNPs on an *ad hoc* basis. In contrast, CWAS systematically integrates these readouts in a statistically rigorous framework that assesses cistrome-wide significance. Importantly, measuring both allelic imbalance and cQTLs increases the number of peaks under detectable genetic control by roughly 50%. This likely reflects distinctive properties of these two methods: AI measurements achieve high sensitivity by using intra-individual comparisons of allele counts that control for effects not mediated by cis-SNP effects. On the other hand, AI detection requires heterozygous SNPs across multiple individuals, while cQTL analyses do not.

CWAS illuminates a critical blind spot of TWAS/eQTL-based approaches: genes with complex regulation and context-dependent expression^28, 44^. These genes were depleted for genetic models of expression based on *cis*-SNPs, but contained the most nearby genetic models of AR binding or regulatory element activity. Strikingly, CWAS identified epigenome-trait association in the absence of a high-confidence transcriptome-trait (TWAS) association at 29% of prostate cancer risk regions. Compared to TWAS+ prostate cancer risk regions, genes in CWAS+/TWAS- regions were subject to more complex regulation and were enriched for transcription factors. This attribute allowed us to implicate key prostate developmental genes and proto-oncogenes in prostate cancer genetics that have largely been overlooked because their expression levels at steady state are highly regulated and have limited *cis*-SNP heritability.

We hypothesize that cQTLs in CWAS+/TWAS-prostate cancer risk regions are context-dependent eQTLs. These variants may affect gene expression in specific tissues or cellular conditions that are relevant to prostate cancer, but their individual effects are obscured at steady state. The *NKX3-1* enhancer provides an example. Mutation of rs1160267 – a cQTL within the enhancer – modestly affects NKX3-1 expression at baseline, but this effect is amplified substantially with androgen stimulation^60^. Context-dependent eQTLs frequently alter chromatin “priming” in the absence of stimuli required to elicit effects on gene expression^41^, potentially explaining how the effects of these variants are visible in steady-state chromatin. Our androgen stimulation experiments provides additional evidence of this phenomenon. Transcripts with androgen-induced allelic imbalance tend to harbor nearby regulatory elements that are already imbalanced in the absence of stimulation. By identifying genetic effects on context-specific gene regulation, CWAS may bypass the need to perform eQTL analyses across many conditions to elicit latent eQTLs.

CWAS stands out from other emerging approaches for annotating risk SNPs. The effect of genetic variation on TF binding was recently assessed *in vitro* on an unprecedented scale by assaying 95,886 SNPs with SNP-SELEX^83^. This effort identified a median of 53 SNPs with allelic imbalance for a given TF, compared to 5,580 genetically determined ARBS identified by our approach. CWAS achieves this increase of two orders of magnitude by assessing AR binding at millions of SNPs *in vivo*. In addition, CWAS can capture effects of SNPs on cooperative binding that are not evident when assaying individual TFs *in vitro*. CWAS also contributes orthogonal information from growing databases of reference epigenomes. A recent study of 10,000 epigenomes across 800 tissue samples annotated GWAS risk SNPs for hundreds of traits based on proximity to tissue-specific enhancers^84^. Only one CWAS ARBS out of 74 overlapped with enhancers putatively linked to GWAS risk by this study, highlighting the importance of assaying disease-relevant transcription factors in addition to histone marks^43^.

Furthermore, our approach captures mechanisms that will be missed by the common practice of intersecting epigenomic features on GWAS risk signal: the top cQTLs for 50% of ARBS cQTLs and 65% of H3K27ac cQTLs lie more than 10kb from the peak center. In some cases, the top cQTL may simply tag the causal variant, but the finding that 16% of ARBS cPeaks contain no SNPs indicates that at least a minority are genetically determined by distal *cis*-SNPs. Ultimately, applying CWAS to growing epigenomic reference panels may increase their power for GWAS annotation substantially.

Our approach has several limitations. First, epigenomic peak intensity may correlate with, but not mediate risk. While this concern has been described for TWAS^10^, it is less likely to confound CWAS studies. This is because epigenomic features can often be linked mechanistically to a single nearby variant, as we observed for hundreds of cQTL SNPs that disrupt a TF binding motif within an ARBS. In these cases, the most parsimonious explanation is that the genetic variant directly alters TF binding, which mediates risk. Nonetheless, pleiotropic effects of variants that alter TF binding but affect risk through an independent mechanism are plausible and future studies will be required to determine their prevalence. A second limitation is that CWAS cannot distinguish the contributions of multiple TFs that cooperatively bind at a site – for instance AR, HOXB13, and FOXA1. Many CWAS ARBS may act on co-binding TFs rather than AR, since the cognate motifs of AR cofactors were often disrupted at allelically imbalanced ARBS. An additional limitation is that epigenomic reference panels from many individuals do not yet exist for most tissues and TFs, especially for populations of non-European ancestry. Nevertheless, our results demonstrate that CWAS is powered to detect associations with smaller sample sizes than TWAS or eQTL studies. In addition, our approach for imputing genotypes form epigenomic data in the absence of genotyping will enable analysis of a large amount of existing public data. Ongoing efforts to perform epigenomic profiling on genetically diverse tissues will advance the utility of this approach further.

The strategy we describe charts a path for future analyses to uncover mechanistic insights into the thousands of variant-trait associations that may not colocalize with steady-state gene expression. While we focused on prostate cancer and AR, CWAS can be applied in a vast range of contexts. Because transcriptional biology often underlies complex phenotypes, CWAS should be a powerful and generalizable approach to ascertaining mechanisms of trait and disease heritability. Using our method to infer accurate genotypes from ChIP-seq data, this approach can be applied to existing and future ChIP-seq datasets across any trait for which GWAS summary statistics are available. Importantly, the increased power for discovery afforded by CWAS unlocks the ability to study the genetics of human disease in smaller populations of particular interest, such as patients enrolled in clinical trials.

## Supplemental Figures

**Figure S1.**
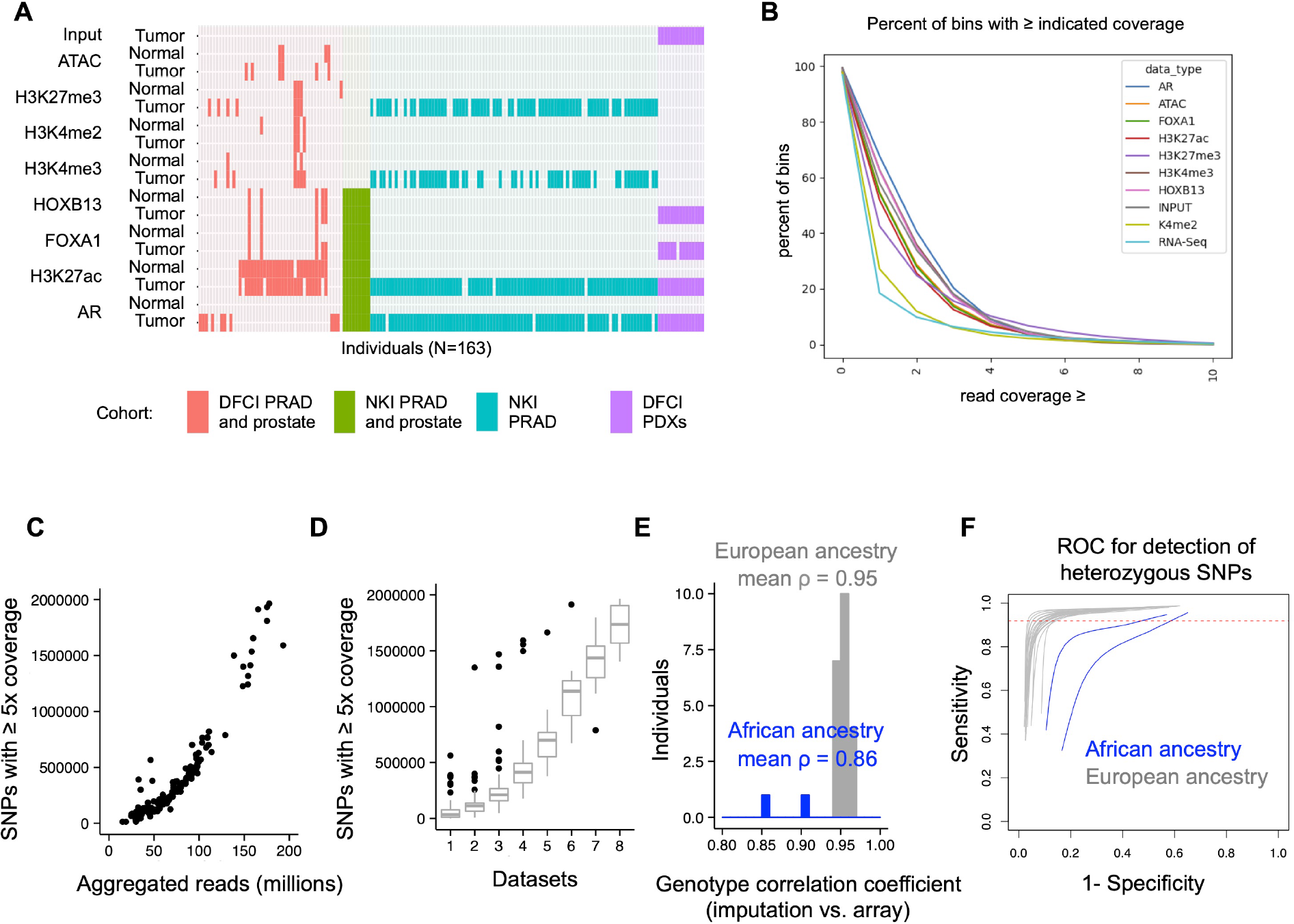
Accurate genotyping of SNPs from epigenomic data. (**A**) Overview of 575 epigenomic datasets merged across 163 individuals for genotyping. Datasets are colored by cohort (See Table S1). (**B**) Genomic distribution of reads in ChIP-seq, RNA-seq and input control (whole genome) data. The genome was divided into non-overlapping 500 base-pair windows and cumulative read counts for each bin were summed. For each datatype, five samples were randomly selected and down-sampled to 8.4 million reads for uniformity. The mean percentage of bins with the indicated number of read counts is shown for each datatype. (**C**) Number of covered SNPs (≥ 5 reads) versus total aggregated reads for each individual. (**D**) Number of covered SNPs (≥ 5 reads) for each individual as the indicated number of datasets are merged. Datasets were added in random order for a given individual. (**E**) Correlation of imputed versus array-based genotype dosages across 24 individuals. (**F**) Receiver operating characteristic curve for detection of heterozygous SNPs using sequencing and imputation, with array-based genotypes as ground truth. Dotted red line indicates a mean sensitivity of 0.92 at a specificity of 0.9 in individuals of European ancestry.

**Figure S2.**
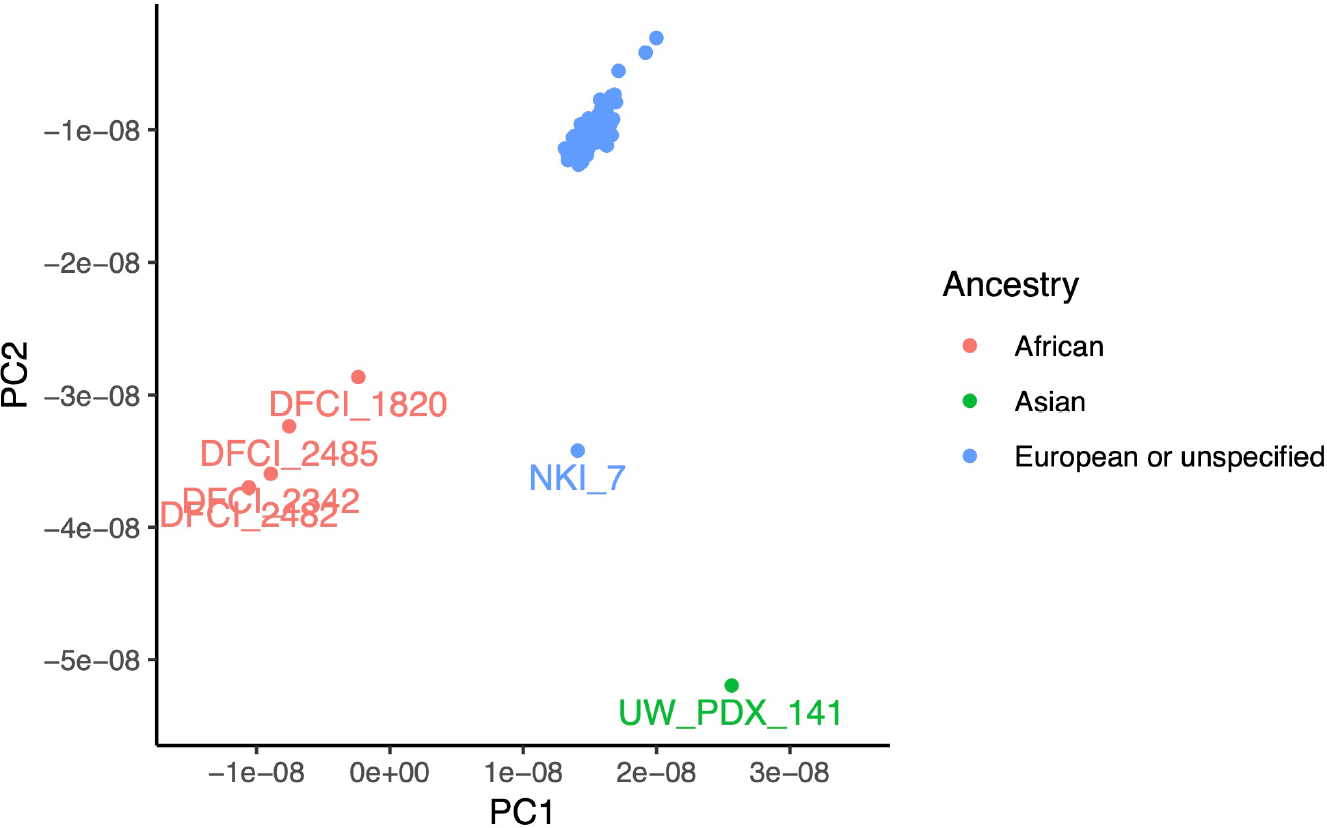
Inferred ancestry of individuals in the study. Projection of imputed genotypes onto the first two principal components of continental ancestry from ref.^86^. Individual identifiers for outlier samples (with values > 2 × standard deviation) are labeled. Self-reported ancestry is coded by color.

**Figure S3.**
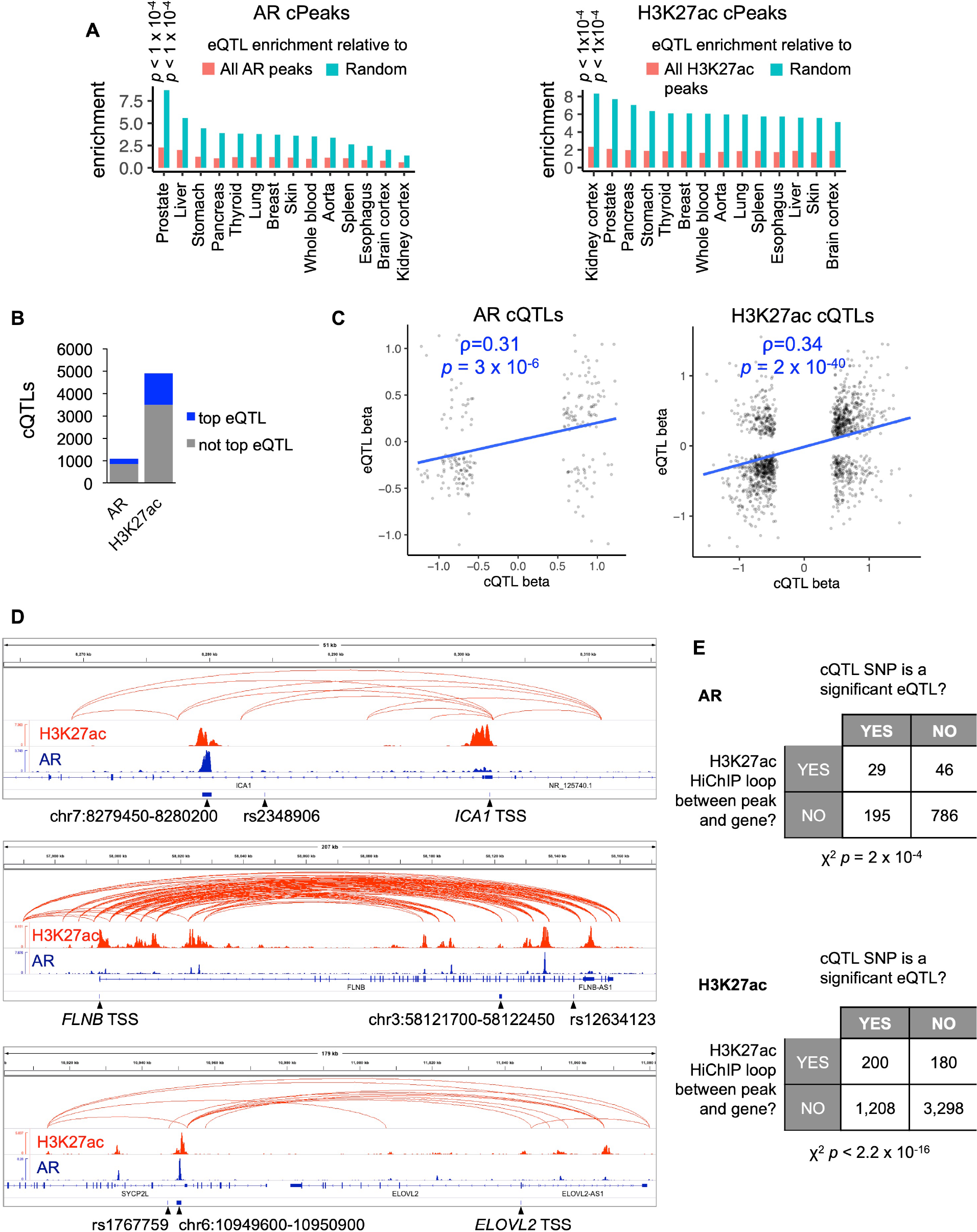
Overlap of cQTLs with prostate tissue eQTLs. (**A**) Enrichment of genetically-determined AR peaks (left) and H3K27ac peaks (right) for overlap with GWAS risk SNPs eQTLs across various tissues. (**B**) number of AR and H3K27ac cQTLs that are also the top eQTL for a gene in prostate tissue. (**C**) correlation of cQTL and eQTL effect size (*β*) for cQTL SNPs. (**D**) Examples of SNPs (labeled with rs identifier) that are both AR cQTLs and eQTLs where the corresponding cPeak and eGene are connected by an H3K27ac HiChIP loop in LNCaP. cPeak coordinates are shown and eGene transcriptional start sites (TSS) is denoted. (**E**) Contingency table showing enrichment of H3K27ac HiChIP looping between the corresponding cPeak and eGene for cQTLs that are also eQTLs.

**Figure S4.**
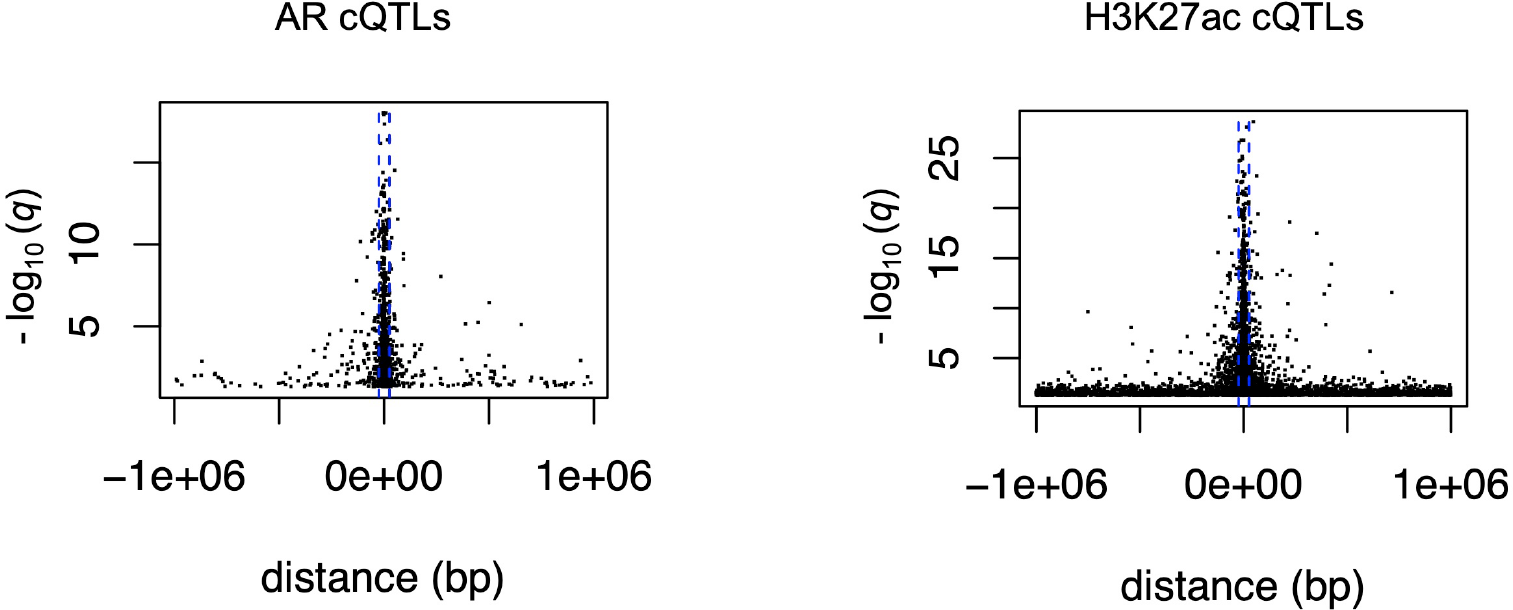
Distribution of cQTLs around cPeaks. cQTL SNP significance versus distance to the center of the corresponding cPeak for significant cQTLs (permutation-based q-value < 0.05). Dashed blue lines indicate ± 25Kb from the peak center.

**Figure S5.**
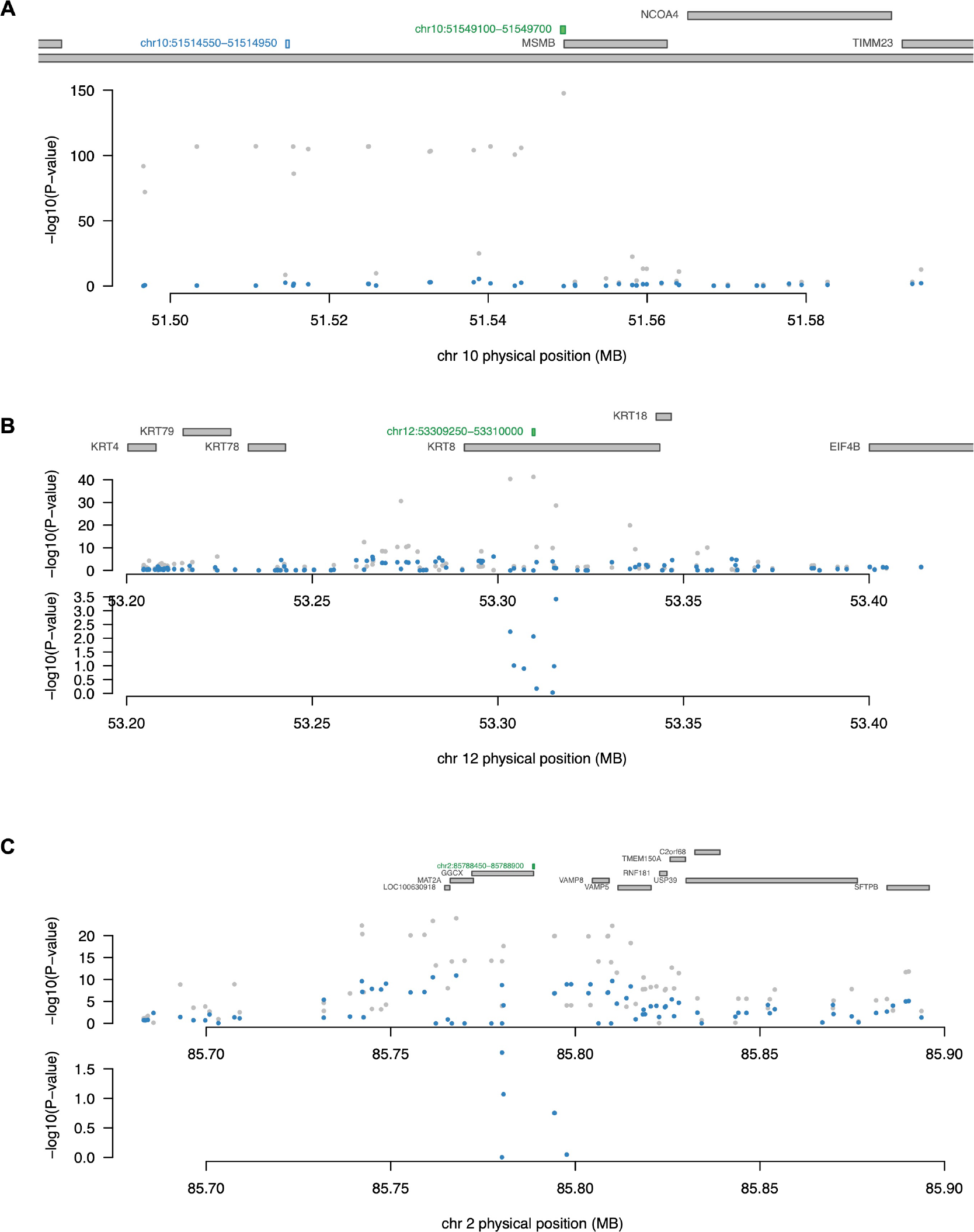
Conditioning of GWAS SNP significance on genetically predicted CWAS AR binding. Genomic context of AR CWAS ARBS (depicted in green) that are significantly associated with prostate cancer risk. Manhattan plots indicate significance of SNP associations with prostate cancer before and after conditioning on genetically predicted CWAS ARBS activity. (**A**) and (**B**) show representative examples where ARBS explain most of the nearby *cis*-SNP GWAS significance. (**C**) CWAS ARBS at the promoter of *GGCX*, where residual GWAS significance remains after conditioning on ARBS, suggesting additional mechanisms underlying risk conferred by SNPs in this region.

**Figure S6.**
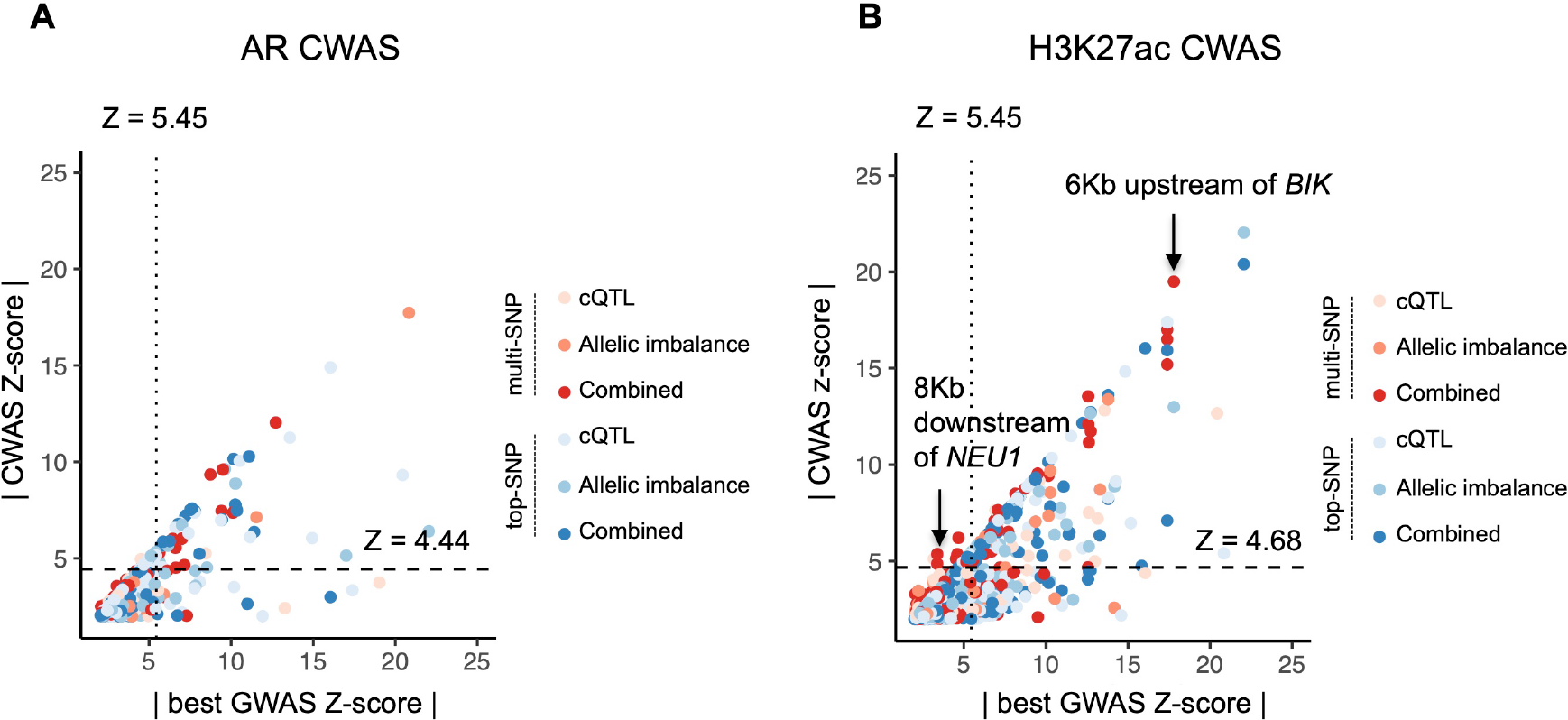
Comparison of CWAS and GWAS significance for tested ARBS and H3K27ac peaks. The absolute value of the association Z-score is plotted for CWAS peak-trait associations (y-axis) and GWAS SNP-trait associations for the most significant nearby SNP (x-axis). (**A**) shows ARBS and (**B**) shows H3K27ac peaks. Dashed horizontal lines indicate genome-wide significance thresholds for CWAS. Vertical dotted lines indicate the GWAS significance threshold of *z* = 5.45.

**Figure S7.**
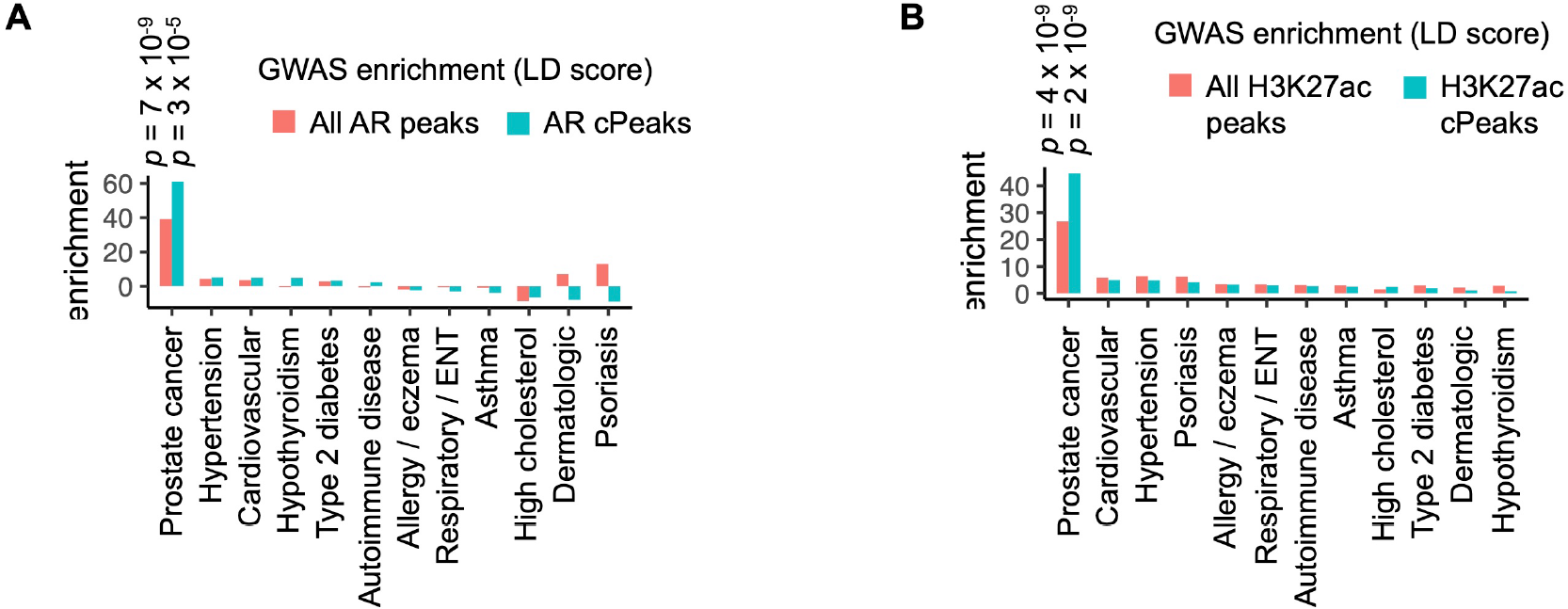
Enrichment of prostate cancer GWAS risk SNPs in genetically determined AR peaks (**A**) and H3K27ac peaks (**B**), as assessed by linkage disequilibrium score regression^6^.

**Fig S8.**
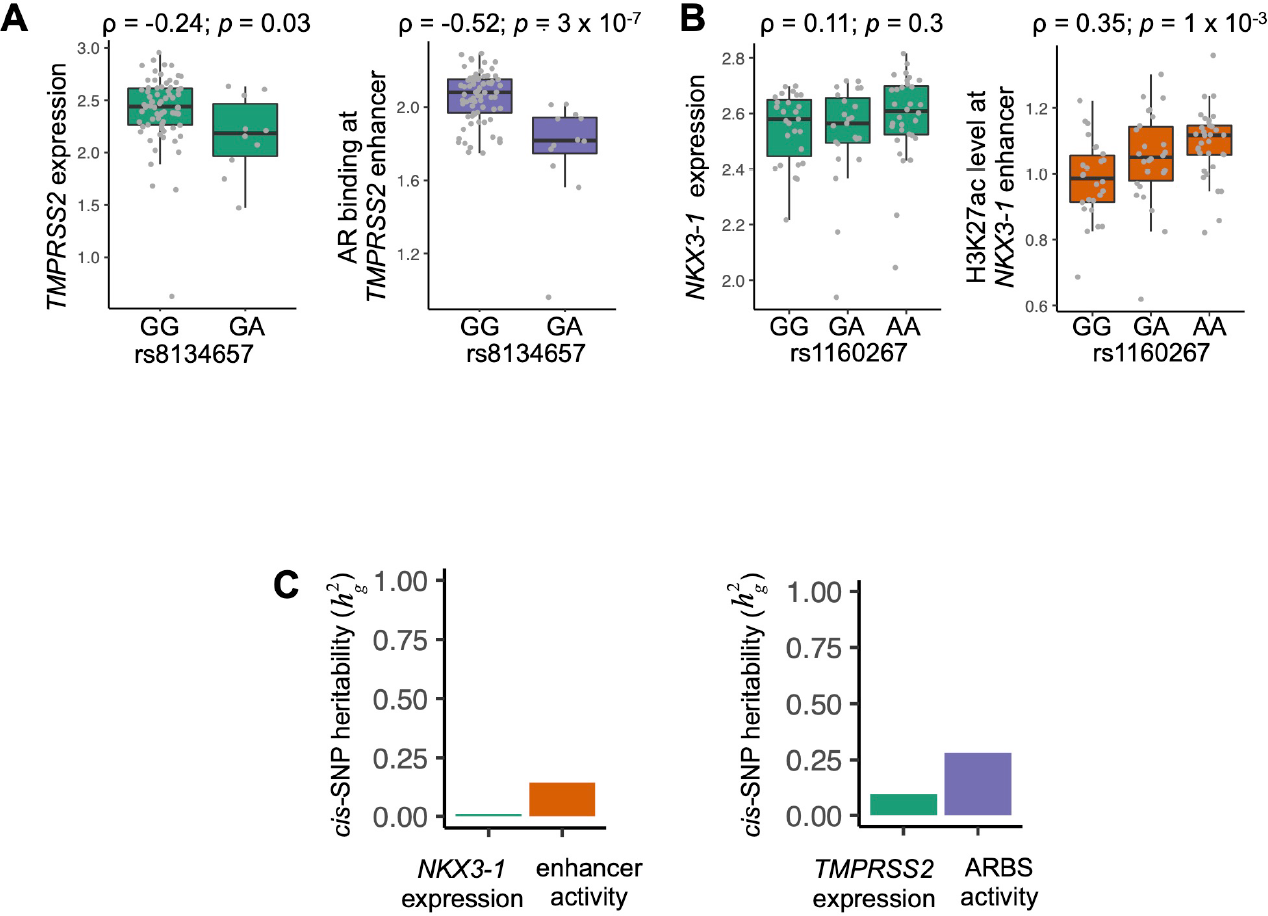
cQTL *vs*. eQTL activity at *TMPRSS2* and *NKX3-1* loci. (**A**) Normalized AR ChIP-seq reads at the *TMPRSS2* enhancer and *TMPRSS2* expression stratified by genotype of the indicated SNP. ρ indicates Pearson correlation coefficient. (**B**) Normalized H3K27ac ChIP-seq reads at the *NKX3-1* enhancer and *NKX3-1* expression stratified by genotype of the indicated SNP. (**C**) Estimated *cis*-SNP heritability for the indicated epigenomic features and corresponding genes.

## Supplemental Tables

**Table S1.** Public datasets used in the study

**Table S2.** Epigenomic dataset quality indicators

**Table S3.** Integrative genetic models of AR binding

**Table S4.** Integrative genetic models of H3K27 acetylation

**Table S5.** qPCR primer and sgRNA sequences

**Table S6.** Significant prostate cancer AR CWAS associations

**Table S7.** Significant prostate cancer H3K27ac CWAS associations

**Table S8.** Significant testosterone level AR CWAS associations

**Table S9.** Significant AR CWAS associations for BPH

**Table S10.** Significant AR CWAS associations for response to androgen deprivation therapy

## Methods

### ChIP-seq peak calling

ChIP-seq fastq files from ref^48^ were downloaded from SRA using SRA toolkit fastq dump v 2.10.0. For uniformity, only the first read in a pair was used for paired-end sequencing datasets. Epigenomic datasets previously generated by our group were processed as described^49, 87^; these data are also available in GEO under accession numbers GSE130408 and GSE161948. ChIP-seq reads were aligned to the human genome build hg19 using the Burrows-Wheeler Aligner (BWA) version 0.7.17^88^. Non-uniquely mapping and duplicate reads were discarded. MACS v2.1.1.20140616^89^ was used for ChIP-seq peak calling with a q-value (FDR) threshold of 0.01. ChIP-seq data quality was evaluated by a variety of measures, including total peak number, FrIP (fraction of reads in peak) score, number of high-confidence peaks (enriched > ten-fold over background), and percent of peak overlap with DHS peaks derived form the ENCODE project. IGV v2.8.2^90^ was used to visualize normalized ChIP-seq read counts at specific genomic loci. Overlap of ChIP-seq peaks and genomic intervals was assessed using BEDTools v2.26.0. Peaks were considered overlapping if they shared one or more base-pairs. Fisher’s test for overlap was performed using the BEDTools fisher command.

### Genotype imputation

We imputed genotypes at 5,495,776 autosomal SNPs present at minor allele frequency > 5% in the Haplotype Reference Consortium (HRC) v1.191^50^. Bam files from epigenomic datasets were merged for each individual using SAMtools merge and run through STITCH v1.6.2^47^ with the following parameters: k=10, ngen=1240, niterations=40, method=diploid (https://hub.docker.com/r/stefangroha/stitch_gcs/tags). The imputation reference panel contained haplotypes of 2,505 individuals in the 1000 Genomes Project Phase 3^85^.

To ensure that individual bam files were correctly assigned to an individual, we used the mpileup and call functions from bcftools v1.9 to call genotypes at 100,000 SNPs and bcftools gtcheck function to test pairwise correlation of homozygous SNP across all files. Samples were clustered based on correlation. Six bam files out of 581 that clustered in a cluster of a different individual were excluded from the analysis.

24 samples were subject to genotyping with Infinium Global Screening Array-24, version 1.0 (Illumina) at the Broad Institute Genomic Services, Cambridge, MA. The Pearson correlation coefficient of allele dosages between imputed and array-based genotypes was evaluated using the R function cor(). A receiver operating characteristic curve was constructed comparing the true positive fraction vs. false positive fraction across cutoffs for genotype dosages.

### Ancestry inference

Ancestry was inferred from imputed SNP genotypes. Samples were projected along the first two principal components of SNP genotypes from reference samples, corresponding to continental ancestry^86^. Projections were computed with plink2 --score no-mean-imputation. Self-reported ancestry was annotated where available.

### Genetic models of epigenomic features

Total and allele-specific peak intensity for H3K27ac and AR were modeled based on *cis*-SNP genotypes in the following steps, which are incorporated into a Snakemake^91^ workflow available at https://github.com/scbaca/cwas.

#### Consensus peak calling

We create a consensus set of H3K27ac and AR by dividing the genome into 50bp windows and including any window with peaks in > 5% of samples. Windows were buffered by 100bp and merged to create a set of 48,948 AR peaks and 81,150 H3K27ac peaks.

#### Allelic imbalance analysis

ChIP-seq reads were analyzed for imbalance of heterozygous SNP alleles using stratAS^42^ (https://github.com/gusevlab/stratAS). Several upstream steps were performed to boost power and accuracy of allelic imbalance detection. Imputed SNP genotypes were phased with Eagle2^92^ using the Sanger Imputation Service (https://imputation.sanger.ac.uk/). Heterozygous SNPs were filtered for mapping bias via the WASP pipeline^93^ and allele-specific read counts were tabulated using ASEReadCounter from the Genome Analysis Toolkit v3.8103^94^.

Briefly, stratAS identifies allelic imbalance by modeling the reads from heterozygous SNPs with a beta-binomial distribution. At each ChIP-seq peak, stratAS takes advantage of haplotype phasing to sum read counts from nearby heterozygous SNP alleles on the same haplotype for each individual. stratAS models the reads from individual *i* overlapping heterozygous germline SNP *j* as: *R_alt,i_* | *R_ref,i_ BetaBin*(*π_j_, ρ_ij_*), where *π* is the mean allelic ratio and *ρ* is a locally-defined, per-individual sequence read correlation parameter reflecting over-dispersion.

Copy number profiles were estimated from off-target ChIP-seq reads with CopywriteR^95^ and used in the modeling of the over-dispersion parameter *ρ*, in order to account for over-dispersion in regions of cancer-associated copy number alterations. *ρ* is estimated for each individual from all heterozygous read-carrying SNPs across ten declines of estimated copy number levels stratAS params.R script, with the following options: --min_snps 50, --min_cov 5, --group 10.

We tested variants with ≥ 20 informative reads within consensus AR and H3K27ac peaks defined above for imbalance. The following additional parameters were set for the stratas.R script: --max_rho 0.2, --window -1, min_cov 1, and --fill_cnv TRUE.

Allelelic imbalance p-values were FDR-adjusted with the qvalue R package (v2.18). Peaks were considered significantly imbalanced if they contained one or more SNPs with imbalance at q < 0.05.

#### Imbalanced SNPs in TF binding motifs

Homer v4.10 was used to identify the most significantly enriched motifs *de novo* among a random selection of 10,000 AR consensus peaks. Imbalanced heterozygous SNPs were tested for overlap with one of these motifs for either allele. Where heterozygous SNPs overlapped, the difference in PWM score between reference and alternate alleles was compared to the allele fraction of reference vs. alternate alleles.

#### cQTL detection

QTLtools v1.2^96^ was used for cQTL detection. Rpkm for each sample at AR and H3K27ac consensus peaks was calculated for each bam file using QTLtools quan with the following flags: --filter-mismatch 5 --filter-mismatch-total 5 --filter-mapping-quality 30. Peaks with a summed rpkm < 10 across all samples were discarded. A covariate matrix was constructed using QTLtools pca --scale --center. Permutation-based p-values^96^ for SNP-peak pairs within a 1Mb window were assessed for cQTLs with QTLtools cis (--normal --permute 1000) after regressing out the first 6 principal components of the peak rpkm covariate matrix. We plotted the distribution of distances between these cPeak-cQTL pairs. After finding that the majority of cQTLs SNPs were within 25kb of the corresponding peak, we also took a focused approach and calculated nominal p-values for *cis*-snp pairs within 25kb, forgoing permutation, which was often not possible for at a distance of 25kb due to a limited numbers of peaks for permutation. These p-values were adjusted by FDR correction and included in downstream analysis where q < 0.05.

For peaks that were tested for both allelic imbalance and cQTLs, combined significance was assessed by combining p-values from the two tests combined using Stouffer’s method^51, 97^.

#### cQTL peak enrichment analysis

Enrichment of eQTL SNPs in cPeaks was tested by permutation. We counted the number of eQTLs for each tissue type overlapping AR or H3K27ac cPeaks and divided this number by the total base-pairs covered by these peaks. We then performed this process on 5,000 same-sized samplings of the complete set of AR or H3K27ac peaks to generate a null distribution. We reported the ratio of peak territory containing a cQTL SNPs in the observed versus simulated data to calculate enrichment and a one-sided p-value. We also calculated enrichment compared to random background by repeating this process using random intervals matched to cPeaks for size, number, and chromosome.

### CWAS model construction

Conventional TWAS models train a predictor of gene expression. Here we extended these models to additionally incorporate allele-specific information and a chromatin phenotype (similar to recent models proposed in the context of statistical fine-mapping^25^ and gene expression^52^). For a given chromatin peak, we take as input the following: a vector of total chromatin activity *y_total_*, with each row containing an individual; the vector of allelic chromatin activity *y_allelic_*, defined as log (*N_p_*/*N_m_*) where *N*_*_ is the total number of reads mapping to the heterozygous variants of the maternal/paternal haplotype, and undefined otherwise; and the matrices of phased maternal and paternal haplotypes ***H***_*p*_ and ***H**_m_*, with individuals as rows and variants within the locus window as columns, containing 0/1 indicators for reference or alternative alleles. We note that maternal or paternal haplotypes can be defined arbitrarily as long as the definition is consistent between the phased genotyped and the allelic reads. In model 1 (“cQTL model”), the relationship between total chromatin activity and genotype is modelled *y_total_* ∼ ***X**_total_* + *ε*, where ***X**_total_* = ***H**_p_* + ***H**_m_* and corresponds to the 0/1/2 allelic dosage for each sample and variant. This model is identical to the models used for conventional TWAS prediction. In model 2 (“allelic imbalance model”), following ref^25^ and ref^52^, the relationship between allelic chromatin activity and haplotype is modelled as *y_allelic_* ∼ ***X**_allelic_* + *ε*, where ***X**_allelic_* = ***H**_p_* − ***H**_m_* and corresponds to the -1/0/1 allele phase. Lastly, in model 3 (“combined model”), we define a “combined” model as 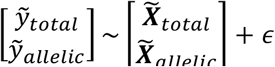 where the twiddle over a variable indicates scaling the columns to zero mean and unit variance. Each model was then fit using LASSO penalized regression to learn genotype to phenotype predictor weights *W* across all variants included in the model (previous work has shown that LASSO models preform comparably to other penalization schemes^98^). Predictive accuracy was evaluated by five-fold cross validation and quantified as the Pearson correlation to the true *y_total_* or *y_allelic_* phenotype. All other model parameters (specifically the LASSO penalty) were fit by nested cross-validation within each training fold.

This analysis is implemented using stratAS with the --predict flag, with --window set to 25kb to include SNPs within 25kb of the peak center.

### Heritability calculations

Heritability of total peak intensity attributable to *cis*-SNPs within 500kb of the peak center 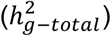 was calculated by the FUSION pipeline as described^45, 53^. Briefly, we model the residual variance of total peak intensity (after centering, scaling, and regressing out the first 6 principal components as fixed-effect covariates) as

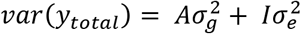

where *A* is the kinship matrix (estimated from cis-SNPs by plink2R v1.1 --make-grm-bin), *I* is the identity matrix, and *σ_g_*^2^ and *σ_e_*^2^ are the variance explained by the *cis*-snps and environment, respectively. The variance parameters are estimated by the restricted maximum likelihood (REML) method with GCTA v 1.26^99^. 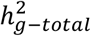 is defined as the ratio of *cis*-snp to total variance in peak intensity 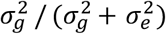. This estimate of 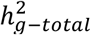 was used to compare cis-snp heritability of AR peaks, H3K27ac peaks and gene expression. In addition, a separate estimate of cis-SNP heritability 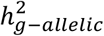 was calculated for the allelic imbalance phenotype, *y_allelic_*, using Haseman-Elston regression^100^. For a given CWAS peak model, 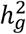 is reported as the estimate 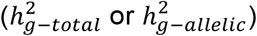 with a smaller p-value.

### CWAS analysis

Integrative models of cQTL and AI were built as described above for each consensus AR or H3K27ac peak based on genotypes of *cis*-SNPs within 25kb (the number of significant models was largely insensitive to the window size, see **Supplemental Note**). We selected the model type with the most significant cross-validation p-value for each peak, and then retained only models with cross validation significance at an FDR of 0.05 across all peaks. The genetic association between predicted peak cQTL activity or AI and GWAS risk was calculated by FUSION, accounting for linkage disequilibrium^45, 53^. FUSION considers the Z-score for genetic peak-trait association as

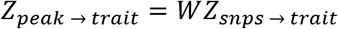

where *Z_snps_*_→*trait*_ is a vector of snp-trait association Z-scores from GWAS summary statistics

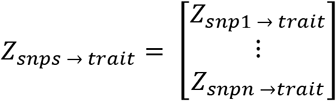

and *W* is a weight matrix defined as

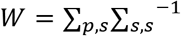

∑*_p_*_,*s*_ is the peak-snp covariance matrix, and ∑*_s_*_,*s*_ is snp-snp covariance matrix, representing linkage disequilibrium. In practice, *W* is learned from the data through penalized regression. Assuming a normal distribution of *Z_peak_*_→*trait*_ around 0, then Z-score for a peak-trait CWAS association is

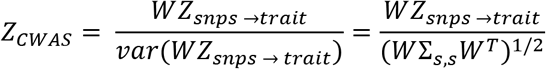

and the corresponding two-sided p-value is obtained from the normal distribution *N*(0,1). CWAS associations were considered significant if p < 0.05 after Bonferonni correction for all peaks of a given type tested (N=5,580 for AR and 17,199 for H3K27ac).

GWAS datasets used in this study are listed in Table S1.

### Conditioning of GWAS SNP significance on CWAS models

FUSION calculated the residual GWAS SNP significance after conditioning on the *cis*-SNP genetic component of CWAS peak cQTL activity or AI as described^101, 102^. Briefly, peaks reaching cistrome-wide significance are grouped into loci if within 100kb of each other. At each locus, a joint model is built to combine peak intensity predictions for the most significant CWAS peaks. CWAS peaks are added iteratively to the joint model from largest to smallest CWAS Z^2^. For each iteration, the CWAS effect size *β* for each peak *i* not in the joint model is conditioned upon the joint model containing peaks *i* +1 to *n*:

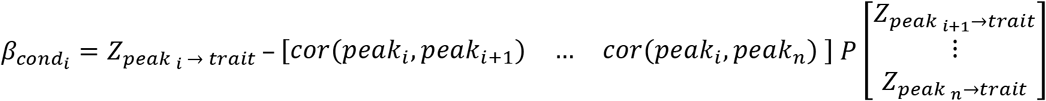

where *P* is a matrix of peak-peak Pearson correlations from cis-SNP-predicted activity for peaks in the joint model imputed into a 1000 Genomes reference panel. The conditioned Z-score *Z_cond,i_* is updated for each peak

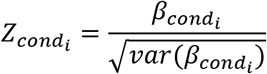

and the process is repeated, adding the peak with largest *Z_cond_* to the joint model until all peaks have been considered or *Z_cond_* for all peaks at the locus is ≤ the 5^th^ percentile of CWAS Z-scores. Peaks are added to the model only if the their correlation r^2^ < 0.3 with all genes already in the model to capture independent peak-trait associations. GWAS SNP associations in the region are then conditioned on the joint CWAS model:

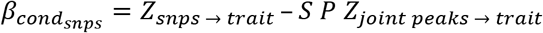

where *S* is a peak-SNP correlation matrix from imputed joint-model peak intensity in 1000 Genomes individuals. Then conditional effect sizes are converted to Z-scores as for peaks. The variance explained for a locus is then calculated as:

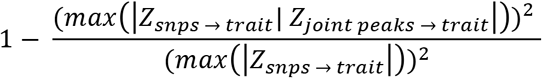

### Overlap of GWAS, TWAS, and CWAS results

Genome-wide significant SNPs (*p* < 5 × 10^-8^) were obtained form published GWAS summary data^56^, assigned hg19 coordinates buffered with 1Mb windows on either side, and merged where windows overlap to obtain 98 prostate cancer GWAS risk regions. Each region was evaluated for overlap with one or more high-confidence CWAS peaks (AR or H3K27ac) or TWAS peaks (from prostate tumor reference panels, or panels incorporating all available tissues). High-confidence peaks and genes were defined as those where 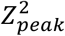 or 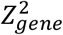 was greater than 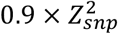 for the most significant GWAS SNP in the region. We elected not to threshold based on statistical colocalization because: (1) no colocalization method currently incorporates allele-specific signal; (2) colocalization methods are highly dependent on the molecular study size and underpowered for hundreds of samples^103^; and (3) colocalization probabilities are highly conservative even in large GWAS^104^. Our high-confidence regions should thus be interpreted as being consistent with explaining the majority of the GWAS variance at the locus.

Prostate cancer risk loci with significant CWAS associations but no significant GWAS associations were evaluated in a large prostate cancer GWAS that was published after this manuscript was prepared^57^. The 269 independent risk variants reported in the ref.^57^ were buffered with 1Mb windows. AR and H3K27ac CWAS peaks were evaluated for overlap with these windows to identify peaks with nearby SNPs that were significant only in the larger GWAS.

### Androgen deprivation therapy GWAS

Men who received androgen deprivation (ADT) for metastatic hormone-sensitive prostate cancer (N=687) were evaluated. 265 of these patients were from the control arm of the CHAARTED clinical trial (E3805)^77^. The remaining 422 were patients treated at Dana-Farber Cancer Institute who gave informed consent for tissue biobanking, genetic analysis, and clinical data collection. These patients were selected to match enrollment criteria for CHAARTED. Subjects were genotyped at approximately one million SNPs with minor allele frequency ≥ 0.05 on Affymetrix 6.0 arrays. Genotypes for SNPs interrogated on the array were called using the Birdsuite algorithm. Alignment to the hg19 genome build was checked using tools provided in SHAPEIT. Strands were flipped using plink when necessary. SHAPEIT was used to pre-phase the SNPs using 1000 Genomes Phase 3 panel as the reference, followed by imputation using IMPUTE (v2.3.1). Time to progression, as assessed in the trial, was evaluated for association with genotypes with the Cox proportional hazards model implemented by the ProbABEL R package^105^. The square root of the corresponding *χ*^2^ statistics were used as the GWAS summary statistics for CWAS analysis.

To limit hypothesis testing, we restricted CWAS association testing to CWAS AR peaks within 1Mb of the top 200 GWAS SNPs by significance (N=789 peaks).

### cQTL-eQTL overlap analysis

Significant cQTLs (FDR-adjuted permutation-based p values < 0.05) were tested for overlap with Prostate GTEx V8 eQTLs. cQTL coordinates were lifted over to hg38 to allow matching to match GTEx eQTLs. The most significant eQTL-eGene and/or cQTL-cPeak association was selected for each SNP. eQTL associations for each SNP were Bonferroni corrected for the number of genes tested for correlation (median N=27). These p-values were then FDR-corrected across 1,058 total tests between each eQTL and its most significantly correlated eGene, and eQTLs with q < 0.05 were retained. cQTLs were annotated as to whether they were also a top eQTL for a gene (**Fig. S4B**) and the Spearman correlation between eQTL and cQTL effect size (*β*) was calculated (**Fig. S4C**)

All permutation-significant cQTLs described above were assessed for their most significant eQTL-eGene association. We tested whether SNPs that are both eQTLs and cQTLs tend to have physically interacting eGenes and ePeaks. We created a contingency table denoting whether each cQTL is also a significant eQTL and whether an H3K27ac HiChIP loop^54^ connects the cPeak with the transcription start site (TSS) of the top eGene. A window of 10kb around loop anchors was used to evaluate TSS or peak overlap with HiChIP anchors.

### DHS-QTL overlap analysis

cQTLs with FDR-adjusted permutation-based q < 0.05 and GTEx prostate eQTLs were tested for overlap with DNAase I hypersensitivity (DHS) peaks from ref^62^. The number of tissues (out of N=733, representing 438 cell types/states) overlapping AR or H3K27ac cQTLs and eQTLs was tabulated. The median number of tissues in which a QTL falls within a DHS peak was plotted and these distributions were compared for AR cQTLs, H3K27ac cQTLs and eQTLs with the Wilcoxon rank-sum test. DHS data mapped to hg19 were used for cQTLs and data mapped to hg38 were used for eQTLs.

### Generation of SNP STARRseq library

SNP STARRseq data were generated for an independent project (unpublished) evaluating allele-specific enhancer activity at prostate cancer GWAS loci. Selected SNPs that were represented in both the SNP STARRseq data and the CWAS analysis are shown in the manuscript.

Pooled human genomic DNA (NA13421; Coriell Institute for Medical Research) was fragmented (500-800bp), end-repaired and ligated with xGen stubby adaptors (IDT) containing 3bp unique molecular identifiers. The target regions were captured using a custom xGen biotinylated oligonucleotide probe pool (IDT) and Dynabeads M-270 Streptavidin beads (IDT). Post-capture was PCR-amplified with STARR_in-fusion_F primer (TAGAGCATGCACCGGACACTCTTTCCCTACACGACGCTCTTCCGATCT) and STARR_in-fusion_R primer (GGCCGAATTCGTCGAGTGACTGGAGTTCAGACGTGTGCTCTTCCGATCT), and then cloned into AgeI-HF (NEB) and SalI-HF (NEB) digested hSTARR-ORI plasmid (Addgene plasmid #99296) with NEBuilder HiFi DNA Assembly Master Mix (NEB). The SNP STARRseq capture library was then transformed into MegaX DH10B T1R electrocompetent cells (Invitrogen) and plasmid DNA was extracted using the Qiagen Plasmid Maxi Kit.

### SNP STARRseq

The SNP STARRseq library (100ug plasmid DNA/replica) was transfected into LNCaP cells (5 × 10^7^ cells/replica; 3 biological replicas) using the Neon Transfection System (Invitrogen). Cells were grown in RPMI 1640 medium supplemented with 10% FBS and collected 48hrs post-electroporation. Cells were lysed with Precellys CKMix Tissue Homogenizing Kit (Bertin Technologies) and total RNA was extracted using Qiagen RNeasy Maxi Kit (Qiagen).

mRNA was isolated with Oligo (dT)25 Dynabeads (Thermo Fisher) and reverse-transcribed with the plasmid-specific primer (CTCATCAATGTATCTTATCATGTCTG). The synthesized SNP STARRseq cDNA was treated with RNaseA and amplified by junction PCR (15 cycles) with the RNA_jPCR_f primer (TCGTGAGGCACTGGGCAG*G*T*G*T*C) and the jPCR_r primer (CTTATCATGTCTGCTCGA*A*G*C). RNAseq was performed with a HiSeq4000 (150bp; PE)

The SNP STARRseq capture library was PCR-amplified with DNA-specific junction PCR primer (DNA_jPCR_f, CCTTTCTCTCCACAGGT*G*T*C) and jPCR_r primer. After purification with Ampure XP beads, DNA was PCR-amplified with TruSeq dual indexing primers (Illumina) to generate Illumina compatible libraries. Libraries were sequenced twice with Illumina MiSeq (75/425 PE for first MiSeq run, and 425/75 PE for second MiSeq run).

### SNP STARRseq analysis

To annotate the STARRseq plasmid library, the two 425 bp reads from the asymmetrical Illumina datasets were matched based on the ∼100 bp of overlapping sequence and then clustered based on the UMIs and insert sequence. The insert consensus sequence was generated from all clusters with >2 reads using Calib^106^ and the bbtools bbmerge function^107^. Bases not matching the consensus sequence were identified and compared to known SNPs in dbSNP. A database was then made to correlate the genomic coordinates of the insert and genetic variants with a 24bp plasmid-specific barcode generated from the insert 5′ and 3′ sequence (3bp UMI + 9 bp of insert sequence at each end). The enhancer activity of each genomic region and variant was then quantified by counting the frequency of the plasmid-specific barcode from the STARRseq mRNA.

To identify SNPs with allelically imbalanced enhancer activity, we conducted differential expression tests to compare expression driven by fragments containing reference vs. alternate alleles for each SNP. Only SNPs with more than three unique plasmids for both reference and alternate alleles were included for analysis. A negative Binomial regression for the enhancer activity was performed for each SNP with the following model using glm.nb() function in MASS R library:

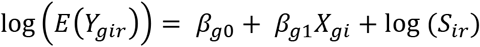

where *Y_gir_* is the RNA read counts of fragment *i* which contains SNP *g* in replicate *r*, *X_gi_* is the genotype of SNP *g* carried by fragment *i* (*X_gi_*=0 for reference allele and *X_gi_*=1 for variant allele), *β_g_*_0_ is the log enhancer activity of reference allele and *β_g_*_1_ is the log fold-change of enhancer activity comparing variant allele versus reference allele, *S_ir_*= *P_i_L_r_* is a fragment and replicate specific normalizing factor, adjusting for the plasmid abundance of fragment *i* (*P_i_*) and replicate library size (*L_r_*).

### CRISPRi suppression of ARBS

The gRNA sequences used to target CWAS enhancers were identified using the CRISPick algorithm (https://portals.broadinstitute.org/gppx/crispick/public). The highest scoring gRNAs near the center of a given peak were selected. The gRNA sequences (Table S5) were synthesized as single stranded oligonucleotides (IDT DNA) with compatible sticky ends (for detailed protocol see https://www.broadinstitute.org/rnai/public/resources/protocols). Annealed oligonucleotides were cloned into lenti_U6sg-KRAB-dCas9-puro using Esp3I. Insert sequences were confirmed by Sanger sequencing performed by the CCR Genomics Core at the National Cancer Institute.

Lentivirus was produced by transfecting 293T cells with the gRNA and KRAB-dCas9 expression plasmid together with the packaging plasmids VsVg (Addgene 12259) and psPax2 (Addgene 12260) using TransIT-LT1 transfection reagent (Mirus). Supernatant containing virus was harvested 48 hours following transfection and used to transduce the LNCaP cell line in the presence of 4 mg/ml polybrene and media exchanged after 24 hours. Conditions were optimized to ensure > 95% transduction as assessed by selection with puromycin. RNA was isolated 4 days after transduction using QIAGEN RNeasy Plus Kit and cDNA synthesized using

NEB Protoscript II First Strand cDNA Synthesis Kit. Quantitative PCR was performed on a Quantstudio 6 using SYBR green. Primers used for qRT-PCR are listed in Table S5. A nontargeting gRNA and gRNA targeting an intergenic region were used as negative controls. Gene expression was normalized to GAPDH and DDCt values were calculated using the nontargeting gRNA as the control sample. Data from three independent biological replicates were used to determine average fold change and data represent the average and standard deviation with significance determined by Student’s t test.

### LNCaP DHT stimulation

LNCaP cells (ATCC CRL-1740) were cultured in phenol red free RPMI (#11835030, Gibco) with 10% charcoal stripped FBS (#100-119, Gembio) for 3 days. then were stimulated with either 10 nM DHT (5α-Androstan-17β-ol-3-one, Dihydrotestosterone, A8380, Sigma) or EtOH (Vehicle) for 16 hours. Subsequently cells were collected for further analysis accordingly. LNCaP cells were authenticated by comparing short tandem repeats to parental LNCaP cells in the ATCC database. Prior to experiments, cells tested for several strains of mycoplasma contamination using LookOut Mycoplasma PCR Detection Kit (Sigma-Aldrich #D9307).

ChIP-seq in LNCaP was performed as previously described^87^ (Ref). Briefly, Ten million cells were fixed with 1 % formaldehyde at room temperature for 10 minutes and quenched with 0.25M glycine, Harvested cells in lysis buffer (1% NP-40, 0.5% sodium deoxycholate, 0.1% SDS and protease inhibitor (#11873580001, Roche) in PBS) were sheared to 300–800 bp chromatin using a Covaris E220 sonicator (140 watt peak incident power, 5% duty cycle, 200 cycleburtst). Sonicated chromatin was subjected to H3K27ac antibody (C15410196, Diagenode) coupled with Dynabeads protein A/G (Life Technology # 10001D, 10003D) overnight at 4 °C. Chromatin was washed in LiCl wash buffer (100 mM Tris pH 7.5, 500 mM LiCl, 1% NP-40, 1% sodium deoxycholate) 6 times for 10 minutes sequentially. Immuno-precipitated chromatin and input were treated with RNase A at 37 °C for 30 minutes and decrosslinked in elution buffer (1% SDS, 0.1 M NaHCO3) with proteinase K for 6–12 hours at 65 °C with gentle rocking. DNA was purified using Qiagen Qiaquick columns (#28104). Libraries were prepared using SMARTer ThruPLEX DNA-Seq Kit (Takara Bio # R400675)

ATAC seq libraries were prepared using Omni-ATAC protocol^108^. Freshly collected 50,000 nuclei in cold lysis buffer (10 mM Tris-HCl, pH 7.5, 10 mM NaCl, 3 mM MgCl2, 0.1% NP-40, 0.1% Tween20, 0.01% Digitonin) were fragmented in 50 µl of transposition mix (25 µl 2× TD buffer, 16.5 µl PBS, 0.5 µl 1% digitonin, 0.5 µl 10% Tween-20, 5 µl water) with 2.5 µl transposase (Illumina 20034197) for 30 min at 37 °C with shaking at 1000 r.p.m. in a thermomixer. DNA was purified using Qiagen MinElute (#28004) and libraries were amplified up to the cycle number determined by 1/3rd maximal qPCR fluorescence

Total mRNA was collected from 300,000 cells using RNA easy kit (Qiagen 74044) with RNase-Free DNase Set (Qiagen no. 79254) according to the manufacturer instructions. RNA purity and concentration were determined on 2100 Bioanalyzer (Agilent) using Agilent RNA 6000 Nano Kit # 5067-1511). 400ng RNA samples were submitted to Novogene for RNA library preparation.

ChIP-seq, RNA-seq, and ATAC-seq libraries and sequenced with 150bp paired-end reads on a HiSeq 250 instrument (Novogene). ChIP-seq and ATAC-seq peaks were called using MACS2 as described above and allelic imbalance in peaks and gene expression was evaluated using stratAS^42^.

## Supplemental Notes

### Accurate genotype imputation from ChIP-seq reads

CWAS requires germline SNP genotypes, which are generally not available for publically available ChIP-seq data. We developed an approach to impute germline genotypes from ChIP-seq reads directly. We merged sequencing reads from 575 epigenomic datasets for the 165 individuals on a per-individual basis (**Fig. S1A**). We compared homozygous SNPs called on each individual data-set at areas of high coverage to verify that merged samples derived from the same individual. Six samples were excluded where genotypes did not correspond closely: (DF_FOXA1_2483_N, DF_FOXA1_2484_N, DF_K4me2_2078_T, P309T_H3K27ac_1, P310T_H3K27ac_1, UW_FOXA1_PDX_189_4). From these merged sequencing reads, we imputed germline genotypes at ∼5.5 million SNPs with a minor allele frequency of 5% or greater in a predominantly European reference population^50^ using STITCH^47^. ChIP-seq data are amenable to this approach because of their broad genomic coverage, compared for example, to RNA-seq data (**Fig. S1B**). By aggregating a median of two datasets per individual (median 70.8 million mapped reads) we achieved ≥5x coverage of 290,149 SNPs (**Fig. S1C**). The addition of up to eight datasets per individual (the maximum available in this study) increased the number of covered SNPs without evidence of saturation (**Fig. S1D**). This finding likely reflects the varying genomic coverage across epigenomic datatypes. Thus, combining epigenomic datasets provides coverage of hundreds of thousands of SNPs, on par with modern genotyping arrays and sufficient for imputation of non-covered SNPs.

We benchmarked imputed genotypes against “ground truth” of genotypes determined by dense SNP arrays for 24 individuals, with a median of two datasets per individual. Imputed genotypes correlated closely with array-based genotypes (Pearson correlation coefficient ρ = 0.95 for individuals of European ancestry and ρ = 0.86 for two individuals of African Ancestry). (**Fig. S1E** and **Fig. S2**). We achieved a mean sensitivity of 0.92 at a specificity of 0.9 for detecting heterozygous SNPs in individuals of European ancestry (**Fig. S1F**), supporting the accuracy of imputation-based genotyping from epigenomic data in the study population.

### Overlap of cQTLs with eQTLs

Several lines of evidence suggest that the genetically regulated peaks we identified drive gene expression. eQTLs were enriched in genetically determined AR and H3K27ac peaks (cPeaks) compared to the full complement of AR and H3K27ac peaks (**Fig. S3A**). 22% of AR cQTLs and 28% of H3K27ac cQTLs were also the top eQTL for a gene in prostate tissue (**Fig. S3B**). The direction and magnitude of a SNP’s effect on its top corresponding gene (eGene) or cPeak correlated significantly (correlation *p* = 7 × 10^-6^ and *p* = 2 × 10^-41^ for AR and H3K27ac cQTLs; **Fig. S3C.**). Peaks and genes that physically interact, as assessed by H3K27ac HiChIP in a prostate cancer cell line^54^, frequently shared the same SNP as a cQTL/eQTL. For 50% (230/457) of cases where a cPeak was connected to a gene promoter by H3K27ac HiChIP loops, the cQTL SNP was also an eQTL for the gene (**Fig. S3D-E**). Thus, *cis*-SNPs that determine AR binding and regulatory element activity also affect expression of physically interacting genes.

### Additional notes epigenomic model generation

We tested multiple window distances around peak centers for modeling epigenomic features. For AR peak models, incorporating SNPs within windows of ± 10kb, 25kb, 50kb, and 100kb resulted respectively in 9,396, 9,679, 9465, and 9,712 peaks with nominal cross-validation significance. These comparable numbers likely reflect the fact that the most significant genetic determinants of peaks tend to fall near the peak center (**Fig. S4**). Given this finding, we chose a window size of ± 25kb, reasoning that although larger windows would better model peaks with more distal genetic determinants, a more narrow peak window would enrich for true positive variants-peak correlations. Consistent with this hypothesis, a ± 25Kb window contains only 69% of all AR associations, but 84% of the top fifth percentile of associations by significance. Similarly, for H3K27ac, a ± 25Kb window contains only 48% of all cQTLs, but 81% of the top fifth percentile of associations by significance. cQTL q-values were significantly smaller within 25kb compared to beyond 25kb of the peak center (*p* = 4 × 10^-21^ and *p* = 4 × 10^-228^ for AR and H3K27ac, respectively).

We considered several schemes for accounting for multiple models (up to 6 model types) tested for each peak. TWAS studies have typically accepted nominally significant cross-validation p-values for gene expression models without correcting for multiple hypothesis testing^53^. To increase stringency, we performed FDR correction after selecting the model type with the most significant cross validation for each peak (N=18,558 for AR and N=49,058 for H3K27ac). The significance of peak-trait associations is subsequently assessed conservatively by Bonferroni correction.

## Author contributions

S.C.B, S.A.G, and S.C.B conceived of the study. S.C.B analyzed the data and wrote the manuscript under the joint supervision of M.L.F. and S.A.G. J.-H. S. generated and C.K. analyzed allelic imbalance data from LNCaP. T.M. and Y.D. analyzed SNP-STARR-seq data under the supervision of N.L. and B.P. C.S. performed CRISPRi experiments under the supervision of D.Y.T. S. Z. assisted with analysis of ChIP-seq data. S.G. assisted with genotype imputation. S. L. and W. Z provided prostate cancer ChIP-seq data. M.M.P. and V.W. analyzed ADT GWAS data preformed on samples and data provided by C.J.S.

## Acknowledgements

S.C.B is supported by grants from the PhRMA Foundation and the Kure It Cancer Research foundation.

## Competing interest

The authors have no competing interests to disclose.

## Data availability

Public datasets used in this study are listed in Table S1. ChIP-seq data generated for this study will be released on the GEO server prior to publication.

## Code availability

Scripts to reproduce the analysis from this study are available at https://github.com/scbaca/cwas.

## Notes

### Competing Interest Statement

The authors have declared no competing interest.

## References

1. https://www.ebi.ac.uk/gwas/.

2. Maurano MT, Humbert R, Rynes E, et al. Systematic localization of common disease-associated variation in regulatory DNA. Science. 2012;337(6099):1190–1195. doi:10.1126/science.1222794

3. Trynka G, Sandor C, Han B, et al. Chromatin marks identify critical cell types for fine mapping complex trait variants. Nat Genet. 2013;45(2):124–130. doi:10.1038/ng.2504

4. Pickrell JK. Joint Analysis of Functional Genomic Data and Genome-wide Association Studies of 18 Human Traits. Am J Hum Genet. 2014;94(4):559–573. doi:10.1016/j.ajhg.2014.03.004

5. Welter D, MacArthur J, Morales J, et al. The NHGRI GWAS Catalog, a curated resource of SNP-trait associations. Nucleic Acids Res. 2014;42(Database issue):D1001–D1006. doi:10.1093/nar/gkt1229

6. Finucane HK, Bulik-Sullivan B, Gusev A, et al. Partitioning heritability by functional annotation using genome-wide association summary statistics. Nat Genet. 2015;47(11):1228–1235. doi:10.1038/ng.3404

7. Gusev A, Lee SH, Trynka G, et al. Partitioning heritability of regulatory and cell-type-specific variants across 11 common diseases. Am J Hum Genet. 2014;95(5):535–552. doi:10.1016/j.ajhg.2014.10.004

8. Hormozdiari F, Gazal S, van de Geijn B, et al. Leveraging molecular quantitative trait loci to understand the genetic architecture of diseases and complex traits. Nat Genet. 2018;50(7):1041–1047. doi:10.1038/s41588-018-0148-2

9. Gallagher MD, Chen-Plotkin AS. The Post-GWAS Era: From Association to Function. Am J Hum Genet. 2018;102(5):717–730. doi:10.1016/j.ajhg.2018.04.002

10. Wainberg M, Sinnott-Armstrong N, Mancuso N, et al. Opportunities and challenges for transcriptome-wide association studies. Nature Genetics. 2019;51(4):592–599. doi:10.1038/s41588-019-0385-z

11. Boyle EA, Li YI, Pritchard JK. An Expanded View of Complex Traits: From Polygenic to Omnigenic. Cell. 2017;169(7):1177–1186. doi:10.1016/j.cell.2017.05.038

12. Wray NR, Wijmenga C, Sullivan PF, Yang J, Visscher PM. Common Disease Is More Complex Than Implied by the Core Gene Omnigenic Model. Cell. 2018;173(7):1573–1580. doi:10.1016/j.cell.2018.05.051

13. Consortium TGte. The Genotype-Tissue Expression (GTEx) pilot analysis: Multitissue gene regulation in humans. Science. 2015;348(6235):648–660. doi:10.1126/science.1262110

14. GTEx Consortium, Laboratory, Data Analysis &Coordinating Center (LDACC)—Analysis Working Group, Statistical Methods groups—Analysis Working Group, et al. Genetic effects on gene expression across human tissues. Nature. 2017;550(7675):204–213. doi:10.1038/nature24277

15. GTEx Consortium. The GTEx Consortium atlas of genetic regulatory effects across human tissues. Science. 2020;369(6509):1318–1330. doi:10.1126/science.aaz1776

16. Kim J, Ghasemzadeh N, Eapen DJ, et al. Gene expression profiles associated with acute myocardial infarction and risk of cardiovascular death. Genome Medicine. 2014;6(5):40. doi:10.1186/gm560

17. Singh T, Levine AP, Smith PJ, Smith AM, Segal AW, Barrett JC. Characterization of expression quantitative trait loci in the human colon. Inflamm Bowel Dis. 2015;21(2):251–256. doi:10.1097/MIB.0000000000000265

18. Ram R, Mehta M, Nguyen QT, et al. Systematic Evaluation Of Genes And Genetic Variants Associated With Type 1 Diabetes Susceptibility. J Immunol. 2016;196(7):3043–3053. doi:10.4049/jimmunol.1502056

19. Gong J, Mei S, Liu C, et al. PancanQTL: systematic identification of cis-eQTLs and trans-eQTLs in 33 cancer types. Nucleic Acids Res. 2018;46(D1):D971–D976. doi:10.1093/nar/gkx861

20. Liu B, Gloudemans MichaelJ, Rao AS, Ingelsson E, Montgomery SB. Abundant associations with gene expression complicate GWAS follow-up. Nat Genet. 2019;51(5):768–769. doi:10.1038/s41588-019-0404-0

21. Strober BJ, Elorbany R, Rhodes K, et al. Dynamic genetic regulation of gene expression during cellular differentiation. Science. 2019;364(6447):1287–1290. doi:10.1126/science.aaw0040

22. Knowles DA, Davis JR, Edgington H, et al. Allele-specific expression reveals interactions between genetic variation and environment. Nat Methods. 2017;14(7):699–702. doi:10.1038/nmeth.4298

23. Ward MC, Banovich NE, Sarkar A, Stephens M, Gilad Y. Dynamic effects of genetic variation on gene expression revealed following hypoxic stress in cardiomyocytes. Elife. 2021;10. doi:10.7554/eLife.57345

24. Kumasaka N, Knights AJ, Gaffney DJ. Fine-mapping cellular QTLs with RASQUAL and ATAC-seq. Nat Genet. 2016;48(2):206–213. doi:10.1038/ng.3467

25. Wang AT, Shetty A, O’Connor E, et al. Allele-Specific QTL Fine Mapping with PLASMA. The American Journal of Human Genetics. 2020;106(2):170–187. doi:10.1016/j.ajhg.2019.12.011

26. Kim-Hellmuth S, Bechheim M, Pütz B, et al. Genetic regulatory effects modified by immune activation contribute to autoimmune disease associations. Nat Commun. 2017;8(1):266. doi:10.1038/s41467-017-00366-1

27. Umans BD, Battle A, Gilad Y. Where Are the Disease-Associated eQTLs? Trends in Genetics. 2020;0(0). doi:10.1016/j.tig.2020.08.009

28. Wang X, Goldstein DB. Enhancer Domains Predict Gene Pathogenicity and Inform Gene Discovery in Complex Disease. Am J Hum Genet. 2020;106(2):215–233. doi:10.1016/j.ajhg.2020.01.012

29. Yao DW, O’Connor LJ, Price AL, Gusev A. Quantifying genetic effects on disease mediated by assayed gene expression levels. Nature Genetics. 2020;52(6):626–633. doi:10.1038/s41588-020-0625-2

30. Chun S, Casparino A, Patsopoulos NA, et al. Limited statistical evidence for shared genetic effects of eQTLs and autoimmune-disease-associated loci in three major immune-cell types. Nat Genet. 2017;49(4):600–605. doi:10.1038/ng.3795

31. Li YI, van de Geijn B, Raj A, et al. RNA splicing is a primary link between genetic variation and disease. Science. 2016;352(6285):600–604. doi:10.1126/science.aad9417

32. McVicker G, van de Geijn B, Degner JF, et al. Identification of genetic variants that affect histone modifications in human cells. Science. 2013;342(6159):747–749. doi:10.1126/science.1242429

33. Chen L, Ge B, Casale FP, et al. Genetic Drivers of Epigenetic and Transcriptional Variation in Human Immune Cells. Cell. 2016;167(5):1398–1414.e24. doi:10.1016/j.cell.2016.10.026

34. Waszak SM, Delaneau O, Gschwind AR, et al. Population Variation and Genetic Control of Modular Chromatin Architecture in Humans. Cell. 2015;162(5):1039–1050. doi:10.1016/j.cell.2015.08.001

35. del Rosario RC-H, Poschmann J, Rouam SL, et al. Sensitive detection of chromatin-altering polymorphisms reveals autoimmune disease mechanisms. Nat Methods. 2015;12(5):458–464. doi:10.1038/nmeth.3326

36. Grubert F, Zaugg JB, Kasowski M, et al. Genetic Control of Chromatin States in Humans Involves Local and Distal Chromosomal Interactions. Cell. 2015;162(5):1051–1065. doi:10.1016/j.cell.2015.07.048

37. Gate RE, Cheng CS, Aiden AP, et al. Genetic determinants of co-accessible chromatin regions in activated T cells across humans. Nat Genet. 2018;50(8):1140–1150. doi:10.1038/s41588-018-0156-2

38. Degner JF, Pai AA, Pique-Regi R, et al. DNase I sensitivity QTLs are a major determinant of human expression variation. Nature. 2012;482(7385):390–394. doi:10.1038/nature10808

39. Maurano MT, Haugen E, Sandstrom R, et al. Large-scale identification of sequence variants influencing human transcription factor occupancy in vivo. Nature Genetics. 2015;47(12):1393–1401. doi:10.1038/ng.3432

40. Deplancke B, Alpern D, Gardeux V. The Genetics of Transcription Factor DNA Binding Variation. Cell. 2016;166(3):538–554. doi:10.1016/j.cell.2016.07.012

41. Alasoo K, Rodrigues J, Mukhopadhyay S, et al. Shared genetic effects on chromatin and gene expression indicate a role for enhancer priming in immune response. Nat Genet. 2018;50(3):424–431. doi:10.1038/s41588-018-0046-7

42. Gusev A, Spisak S, Fay AP, et al. Allelic imbalance reveals widespread germline-somatic regulatory differences and prioritizes risk loci in Renal Cell Carcinoma. bioRxiv. Published online May 8, 2019:631150. doi:10.1101/631150

43. Benaglio P, D’Antonio-Chronowska A, Ma W, et al. Allele-specific NKX2-5 binding underlies multiple genetic associations with human electrocardiographic traits. Nature Genetics. 2019;51(10):1506–1517. doi:10.1038/s41588-019-0499-3

44. Kasowski M, Grubert F, Heffelfinger C, et al. Variation in Transcription Factor Binding Among Humans. Science. 2010;328(5975):232–235. doi:10.1126/science.1183621

45. Gusev A, Ko A, Shi H, et al. Integrative approaches for large-scale transcriptome-wide association studies. Nat Genet. 2016;48(3):245–252. doi:10.1038/ng.3506

46. Jiang X, Finucane HK, Schumacher FR, et al. Shared heritability and functional enrichment across six solid cancers. Nat Commun. 2019;10(1):431. doi:10.1038/s41467-018-08054-4

47. Davies RW, Flint J, Myers S, Mott R. Rapid genotype imputation from sequence without reference panels. Nat Genet. 2016;48(8):965–969. doi:10.1038/ng.3594

48. Stelloo S, Nevedomskaya E, Kim Y, et al. Integrative epigenetic taxonomy of primary prostate cancer. Nat Commun. 2018;9. doi:10.1038/s41467-018-07270-2

49. Pomerantz MM, Qiu X, Zhu Y, et al. Prostate cancer reactivates developmental epigenomic programs during metastatic progression. Nature Genetics. 2020;52(8):790–799. doi:10.1038/s41588-020-0664-8

50. McCarthy S, Das S, Kretzschmar W, et al. A reference panel of 64,976 haplotypes for genotype imputation. Nature Genetics. 2016;48(10):1279–1283. doi:10.1038/ng.3643

51. Stouffer SA, Suchman EA, Devinney LC, Star SA, Williams Jr. RM. The American Soldier: Adjustment during Army Life. (Studies in Social Psychology in World War II), Vol. 1. Princeton Univ. Press; 1949:xii, 599.

52. Liang Y, Aguet F, Barbeira AN, Ardlie K, Im HK. A scalable unified framework of total and allele-specific counts for cis-QTL, fine-mapping, and prediction. Nat Commun. 2021;12(1):1424. doi:10.1038/s41467-021-21592-8

53. Mancuso N, Gayther S, Gusev A, et al. Large-scale transcriptome-wide association study identifies new prostate cancer risk regions. Nature Communications. 2018;9(1):4079. doi:10.1038/s41467-018-06302-1

54. Giambartolomei C, Seo J-H, Schwarz T, et al. H3k27ac-HiChIP in prostate cell lines identifies risk genes for prostate cancer susceptibility. bioRxiv. Published online October 25, 2020:2020.10.23.352351. doi:10.1101/2020.10.23.352351

55. Emami NC, Kachuri L, Meyers TJ, et al. Association of imputed prostate cancer transcriptome with disease risk reveals novel mechanisms. Nature Communications. 2019;10(1):3107. doi:10.1038/s41467-019-10808-7

56. Schumacher FR, Al Olama AA, Berndt SI, et al. Association analyses of more than 140,000 men identify 63 new prostate cancer susceptibility loci. Nature Genetics. 2018;50(7):928–936. doi:10.1038/s41588-018-0142-8

57. Conti DV, Darst BF, Moss LC, et al. Trans-ancestry genome-wide association meta-analysis of prostate cancer identifies new susceptibility loci and informs genetic risk prediction. Nature Genetics. 2021;53(1):65–75. doi:10.1038/s41588-020-00748-0

58. Pomerantz MM, Shrestha Y, Flavin RJ, et al. Analysis of the 10q11 Cancer Risk Locus Implicates MSMB and NCOA4 in Human Prostate Tumorigenesis. PLoS Genet. 2010;6(11). doi:10.1371/journal.pgen.1001204

59. Clinckemalie L, Spans L, Dubois V, et al. Androgen regulation of the TMPRSS2 gene and the effect of a SNP in an androgen response element. Mol Endocrinol. 2013;27(12):2028–2040. doi:10.1210/me.2013-1098

60. Zhang Z, Chng KR, Lingadahalli S, et al. An AR-ERG transcriptional signature defined by long-range chromatin interactomes in prostate cancer cells. Genome Res. 2019;29(2):223–235. doi:10.1101/gr.230243.117

61. Yang J, Manolio TA, Pasquale LR, et al. Genome partitioning of genetic variation for complex traits using common SNPs. Nat Genet. 2011;43(6):519–525. doi:10.1038/ng.823

62. Meuleman W, Muratov A, Rynes E, et al. Index and biological spectrum of human DNase I hypersensitive sites. Nature. 2020;584(7820):244–251. doi:10.1038/s41586-020-2559-3

63. Koh CM, Bieberich CJ, Dang CV, Nelson WG, Yegnasubramanian S, De Marzo AM. MYC and Prostate Cancer. Genes Cancer. 2010;1(6):617–628. doi:10.1177/1947601910379132

64. Zhang B, Ci X, Tao R, et al. Klf5 acetylation regulates luminal differentiation of basal progenitors in prostate development and regeneration. Nature Communications. 2020;11(1):997. doi:10.1038/s41467-020-14737-8

65. Bhatia-Gaur R, Donjacour AA, Sciavolino PJ, et al. Roles for Nkx3.1 in prostate development and cancer. Genes Dev. 1999;13(8):966–977.

66. Drobnjak M, Osman I, Scher HI, Fazzari M, Cordon-Cardo C. Overexpression of cyclin D1 is associated with metastatic prostate cancer to bone. Clin Cancer Res. 2000;6(5):1891–1895.

67. Economides KD, Capecchi MR. Hoxb13 is required for normal differentiation and secretory function of the ventral prostate. Development. 2003;130(10):2061–2069. doi:10.1242/dev.00432

68. Wu D, Sunkel B, Chen Z, et al. Three-tiered role of the pioneer factor GATA2 in promoting androgen-dependent gene expression in prostate cancer. Nucleic Acids Res. 2014;42(6):3607–3622. doi:10.1093/nar/gkt1382

69. Reckelhoff Jane F., Zhang Huimin, Granger Joey P. Testosterone Exacerbates Hypertension and Reduces Pressure-Natriuresis in Male Spontaneously Hypertensive Rats. Hypertension. 1998;31(1):435–439. doi:10.1161/01.HYP.31.1.435

70. Kloner RA, Carson C, Dobs A, Kopecky S, Mohler ER. Testosterone and Cardiovascular Disease. Journal of the American College of Cardiology. 2016;67(5):545–557. doi:10.1016/j.jacc.2015.12.005

71. Sudlow C, Gallacher J, Allen N, et al. UK Biobank: An Open Access Resource for Identifying the Causes of a Wide Range of Complex Diseases of Middle and Old Age. PLOS Medicine. 2015;12(3):e1001779. doi:10.1371/journal.pmed.1001779

72. Kim S-M, Kim J-Y, Choe N-W, et al. Regulation of mouse steroidogenesis by WHISTLE and JMJD1C through histone methylation balance. Nucleic Acids Res. 2010;38(19):6389–6403. doi:10.1093/nar/gkq491

73. Jin G, Sun J, Kim S-T, et al. Genome-wide association study identifies a new locus JMJD1C at 10q21 that may influence serum androgen levels in men. Human Molecular Genetics. 2012;21(23):5222–5228. doi:10.1093/hmg/dds361

74. Levasseur A, St-Jean G, Paquet M, Boerboom D, Boyer A. Targeted Disruption of YAP and TAZ Impairs the Maintenance of the Adrenal Cortex. Endocrinology. 2017;158(11):3738–3753. doi:10.1210/en.2017-00098

75. Hawley JR, Zhou S, Arlidge C, et al. Cis-regulatory Element Hijacking Overshadows Topological Changes in Prostate Cancer. bioRxiv. Published online January 6, 2021:2021.01.05.425333. doi:10.1101/2021.01.05.425333

76. Sáez C, González-Baena AC, Japón MA, et al. Expression of basic fibroblast growth factor and its receptors FGFR1 and FGFR2 in human benign prostatic hyperplasia treated with finasteride. The Prostate. 1999;40(2):83–88. doi:https://doi.org/10.1002/(SICI)1097-0045(19990701)40:2<83::AID-PROS3>3.0.CO;2-N

77. Sweeney CJ, Chen Y-H, Carducci M, et al. Chemohormonal Therapy in Metastatic Hormone-Sensitive Prostate Cancer. New England Journal of Medicine. 2015;373(8):737–746. doi:10.1056/NEJMoa1503747

78. Pomerantz M, Wang XV, Kantoff PW, et al. Genome-wide association study (GWAS) of response to androgen deprivation therapy (ADT) and survival in metastatic prostate cancer (PCa). JCO. 2016;34(15_suppl):1540–1540. doi:10.1200/JCO.2016.34.15_suppl.1540

79. Whitaker HC, Shiong LL, Kay JD, et al. N-acetyl-L-aspartyl-L-glutamate peptidase-like 2 is overexpressed in cancer and promotes a pro-migratory and pro-metastatic phenotype. Oncogene. 2014;33(45):5274–5287. doi:10.1038/onc.2013.464

80. African Ancestry Prostate Cancer GWAS Consortium, Berndt SI, Wang Z, et al. Two susceptibility loci identified for prostate cancer aggressiveness. Nat Commun. 2015;6(1):6889. doi:10.1038/ncomms7889

81. Arachchige PD, Carskadon S, Hu J, et al. Abstract 2012: Recurrent rearrangements of NAALADL2 in prostate, breast, cervical, head and neck and lung squamous cell carcinoma. Cancer Res. 2020;80(16 Supplement):2012–2012. doi:10.1158/1538-7445.AM2020-2012

82. Fulco CP, Nasser J, Jones TR, et al. Activity-by-contact model of enhancer–promoter regulation from thousands of CRISPR perturbations. Nature Genetics. 2019;51(12):1664–1669. doi:10.1038/s41588-019-0538-0

83. Yan J, Qiu Y, Ribeiro dos Santos AM, et al. Systematic analysis of binding of transcription factors to noncoding variants. Nature. Published online January 27, 2021:1–5. doi:10.1038/s41586-021-03211-0

84. Boix CA, James BT, Park YP, Meuleman W, Kellis M. Regulatory genomic circuitry of human disease loci by integrative epigenomics. Nature. Published online February 3, 2021:1–8. doi:10.1038/s41586-020-03145-z

85. Auton A, Abecasis GR, Altshuler DM, et al. A global reference for human genetic variation. Nature. 2015;526(7571):68–74. doi:10.1038/nature15393

86. Chen C-Y, Pollack S, Hunter DJ, Hirschhorn JN, Kraft P, Price AL. Improved ancestry inference using weights from external reference panels. Bioinformatics. 2013;29(11):1399–1406. doi:10.1093/bioinformatics/btt144

87. Baca SC, Takeda DY, Seo J-H, et al. Reprogramming of the FOXA1 cistrome in treatment-emergent neuroendocrine prostate cancer. bioRxiv. Published online October 24, 2020:2020.10.23.350793. doi:10.1101/2020.10.23.350793

88. Langmead B, Trapnell C, Pop M, Salzberg SL. Ultrafast and memory-efficient alignment of short DNA sequences to the human genome. Genome Biology. 2009;10(3):R25. doi:10.1186/gb-2009-10-3-r25

89. Zhang Y, Liu T, Meyer CA, et al. Model-based Analysis of ChIP-Seq (MACS). Genome Biology. 2008;9(9):R137. doi:10.1186/gb-2008-9-9-r137

90. Robinson JT, Thorvaldsdóttir H, Winckler W, et al. Integrative Genomics Viewer. Nat Biotechnol. 2011;29(1):24–26. doi:10.1038/nbt.1754

91. Köster J, Rahmann S. Snakemake—a scalable bioinformatics workflow engine. Bioinformatics. 2012;28(19):2520–2522. doi:10.1093/bioinformatics/bts480

92. Loh P-R, Danecek P, Palamara PF, et al. Reference-based phasing using the Haplotype Reference Consortium panel. Nat Genet. 2016;48(11):1443–1448. doi:10.1038/ng.3679

93. van de Geijn B, McVicker G, Gilad Y, Pritchard JK. WASP: allele-specific software for robust molecular quantitative trait locus discovery. Nat Methods. 2015;12(11):1061–1063. doi:10.1038/nmeth.3582

94. Castel SE, Levy-Moonshine A, Mohammadi P, Banks E, Lappalainen T. Tools and best practices for data processing in allelic expression analysis. Genome Biology. 2015;16(1):195. doi:10.1186/s13059-015-0762-6

95. Kuilman T, Velds A, Kemper K, et al. CopywriteR: DNA copy number detection from off-target sequence data. Genome Biology. 2015;16(1):49. doi:10.1186/s13059-015-0617-1

96. Delaneau O, Ongen H, Brown AA, Fort A, Panousis NI, Dermitzakis ET. A complete tool set for molecular QTL discovery and analysis. Nature Communications. 2017;8(1):15452. doi:10.1038/ncomms15452

97. Whitlock MC. Combining probability from independent tests: the weighted Z-method is superior to Fisher’s approach. Journal of Evolutionary Biology. 2005;18(5):1368–1373. doi:https://doi.org/10.1111/j.1420-9101.2005.00917.x

98. Gusev A, Lawrenson K, Lin X, et al. A transcriptome-wide association study of high grade serous epithelial ovarian cancer identifies novel susceptibility genes and splice variants. Nat Genet. 2019;51(5):815–823. doi:10.1038/s41588-019-0395-x

99. Yang J, Lee SH, Goddard ME, Visscher PM. GCTA: A Tool for Genome-wide Complex Trait Analysis. Am J Hum Genet. 2011;88(1):76–82. doi:10.1016/j.ajhg.2010.11.011

100. Haseman JK, Elston RC. The investigation of linkage between a quantitative trait and a marker locus. Behav Genet. 1972;2(1):3–19. doi:10.1007/BF01066731

101. Yang J, Ferreira T, Morris AP, et al. Conditional and joint multiple-SNP analysis of GWAS summary statistics identifies additional variants influencing complex traits. Nat Genet. 2012;44(4):369–375, S1-3. doi:10.1038/ng.2213

102. Gusev A, Mancuso N, Won H, et al. Transcriptome-wide association study of schizophrenia and chromatin activity yields mechanistic disease insights. Nature Genetics. 2018;50(4):538–548. doi:10.1038/s41588-018-0092-1

103. Hukku A, Pividori M, Luca F, Pique-Regi R, Im HK, Wen X. Probabilistic colocalization of genetic variants from complex and molecular traits: promise and limitations. Am J Hum Genet. 2021;108(1):25–35. doi:10.1016/j.ajhg.2020.11.012

104. Barbeira AN, Bonazzola R, Gamazon ER, et al. Exploiting the GTEx resources to decipher the mechanisms at GWAS loci. Genome Biol. 2021;22. doi:10.1186/s13059-020-02252-4

105. Aulchenko YS, Struchalin MV, van Duijn CM. ProbABEL package for genome-wide association analysis of imputed data. BMC Bioinformatics. 2010;11(1):134. doi:10.1186/1471-2105-11-134

106. Orabi B, Erhan E, McConeghy B, et al. Alignment-free clustering of UMI tagged DNA molecules. Bioinformatics. 2019;35(11):1829–1836. doi:10.1093/bioinformatics/bty888

107. Bushnell B. BBMap: A Fast, Accurate, Splice-Aware Aligner. Lawrence Berkeley National Lab. (LBNL), Berkeley, CA (United States); 2014. Accessed April 27, 2021. https://www.osti.gov/biblio/1241166-bbmap-fast-accurate-splice-aware-aligner

108. Corces MR, Trevino AE, Hamilton EG, et al. An improved ATAC-seq protocol reduces background and enables interrogation of frozen tissues. Nat Methods. 2017;14(10):959–962. doi:10.1038/nmeth.4396

